# Self-*S*-sulfonation in a bacterial persulfide dioxygenase mediates thiol persulfide detoxification

**DOI:** 10.64898/2026.02.20.707092

**Authors:** Julius O. Campeciño, Sofia S. Costa, Brenna J. C. Walsh, Jonathan C. Trinidad, Jyoti Kannoujia, Andrew T. Poore, Casey Van Stappen, Giovanni Gonzalez-Gutierrez, Margarida Archer, José A. Brito, David P. Giedroc

## Abstract

A ubiquitous class of non-heme Fe(II) enzymes, the persulfide dioxygenases (PDOs), provide protection against hydrogen sulfide (H_2_S) poisoning. The PDO in humans is a single-domain enzyme, while bacterial PDOs, such as CstB of *Staphylococcus aureus*, are often fused to a sulfurtransferase (rhodanese) module. Canonical PDOs cleave the S-S bond of glutathione persulfide (GSSH) to produce GSH and sulfite (SO_3_^2–^). In contrast, CstB, via an unknown mechanism, converts two RSSH to thiosulfate (S_2_O_3_^2–^) without the release of sulfite. Six crystallographic structures of *S. aureus* CstB reveal that a Cys-Gly sequence (C201-G202) in a CstB-unique dynamic loop functions as a glutathione mimic, occupying one face of the hemifacial octahedral Fe(II) coordination site. We establish that CstB self-*S*-sulfonates C201 in a thiol persulfide, Fe(II) and O_2_-dependent manner, which is then shuttled to a persulfidated C408 in the rhodanese domain ≈27 Å away via electrostatic steering to generate thiosulfate as the sole oxidation product. Both C201A and C408A CstBs are inactive in O_2_-consumption. Self-*S*-sulfonation ensures rapid clearance of diverse reactive sulfur species under conditions where these species accumulate, permitting *S. aureus* to harness their cytoprotective effects while avoiding cellular toxicity.

---

Hydrogen sulfide (H_2_S) is a physiologically important pro-signaling^1^ molecule that can be both harmful and beneficial to cells^2-5^. At physiological levels, H_2_S functions in redox regulation, protection against oxidative stress, and support of energy metabolism in both eukaryotes and prokaryotes. However, at high concentrations, H_2_S inhibits the electron transport chain by targeting cytochrome *c* oxidase in mitochondria of eukaryotic cells and heme-copper-dependent respiratory oxidases in prokaryotes, inducing the expression of sulfide-resistant oxidases^6-8^. Cells encounter H_2_S from both exogenous and endogenous sources, the latter via the transsulfuration and cysteine degradation pathways^9-11^. High levels of exogenous H_2_S found in wastewater streams have been shown to stimulate plasmid conjugation and antibiotic resistance in microbial communities^12^. Sulfide homeostasis therefore allows bacterial cells, particularly in sulfide-rich environments, to manage the dual but opposing impacts of H_2_S^13^.

The first line of defense against sulfide toxicity is often sulfide:quinone oxidoreductase (SQR), an enzyme that catalyzes the oxidation of H_2_S to sulfane sulfur (S^0^), bound to a low-molecular-weight (LMW) thiol such as glutathione (GSH) to form a LMW thiol persulfide, *e.g.*, glutathione persulfide (GSSH)^14, 15^. These persulfides are further processed reductively^16-18^ or oxidatively^19-21^ depending on the sulfide homeostatic pathway the organism possesses, into less toxic sulfur species, which are eventually effluxed or assimilated. For example, the facultative anaerobe *Enterococcus faecalis* harbors a coenzyme A persulfide reductase (CoAPR) that mitigates the toxic effects of an accumulating coenzyme A persulfide (CoASSH) during sulfide stress by reducing the persulfide to coenzyme A and H_2_S, which is likely further metabolized^16^. In aerobic organisms, the sulfane sulfur of LMW persulfides is oxidized to sulfite, regenerating the LMW thiol species^22^. This reaction is catalyzed by members of the persulfide dioxygenase (PDO) superfamily, which includes the mitochondrial ethylmalonic encephalopathy protein 1 (ETHE1), genetic inactivation of which leads to embryonic lethality in humans^21^.

We previously discovered a sulfide detoxification pathway in *Staphylococcus aureus* strain Newman, encoded by the *cst* (<u>c</u>soR-like <u>s</u>ulfurtransferase) operon thought to prevent the toxic accumulation of reactive sulfur species^23^. The operon is regulated by the per- and polysulfide-sensing repressor, CstR, which binds to the operator-promoter binding site and represses transcription when its two cysteine residues are in the reduced state^23, 24^. Upon accumulation of per- and polysulfides, the attacking thiol of CstR quickly forms per- and polysulfide chains, and tri-, and tetrasulfide interprotomer crosslinks with the other cysteine, inhibiting the ability of CstR to bind to the target operator site^23, 24^. This results in derepression of the operon, allowing cellular accumulation of SQR, CstA, CstB, and a candidate TauE^23-25^. This pathway functions as a sequential H_2_S detoxification process in which SQR first catalyzes the transfer of S^2–^ to a low-molecular-weight (LMW) thiol to form LMW thiol persulfide^26^. CstB then converts this persulfide to thiosulfate through its persulfide dioxygenase and sulfurtransferase activities^19^. CstA further processes thiosulfate into sulfite and a CstA-bound persulfide available for downstream acceptors, exploiting its thiosulfate sulfurtransferase function^27^. Finally, the candidate TauE is anticipated to efflux sulfite from cytoplasm although direct support for this proposal remains lacking in *S. aureus*^28^. The core *cst* operon (*tauE-cstR-cstA-cstB*) is duplicated in many methicillin-resistant strains of *S. aureus*, encoded within the methicillin resistance genomic determinant itself^19^, a finding that takes on added significance in the context of the spread of antibiotic resistance in sulfide-rich wastewater environments^12^.

Here, we present structural and mechanistic studies of *Staphylococcus aureus* strain Newman CstB (CstB), a multifunctional PDO-rhodanese fusion protein (PRF) that consists of a N-terminal ETHE1-like PDO domain, a middle pseudorhodanese domain (RHD), and a C-terminal sulfurtransferase domain (ST, Rhod)^19^. Unlike a standalone PDO, which oxidizes the terminal sulfane of GSSH to sulfite, CstB produces no sulfite, instead exclusively making thiosulfate (TS; S_2_O_3_^2–^) from two mol equivalents of thiol persulfide^19^. This process appears to require both the Fe(II)-containing PDO and the ST active sites. Combining crystallography, spectroscopy and extensive mechanistic analysis, we show here that the CstB-specific loop in the PDO domain positions the Sγ atom of a C201-G202 glutathione mimic 3.4 Å away from the Fe(II) center, which when persulfidated, becomes the substrate for Fe(II)- and O_2_-dependent self *S*-sulfonation. This *S*-sulfonated C201 is then transferred to C408 in the Rhod domain, some 27 Å away, attacked by C408 persulfide with the release of TS. O_2_ consumption by the PDO domain is abolished by the C201A and C408A substitutions revealing tight coupling of the two active sites. The C201-harboring “molecular shuttle” is reminiscent of the mobile loops associated with cysteine persulfide transfer in Fe-S cluster protein biogenesis^29^ and the mobile pantothenate “arm” of CoA persulfide reductases^17^ except that here the enzyme shuttles a *S*-sulfonate moiety, an unprecedented finding.

## Results

### Crystallographic structures of CstB

CstB is a multidomain oligomeric enzyme, consisting of the N-terminal non-heme-Fe(II) persulfide dioxygenase (PDO) domain, a middle pseudorhodanese homology domain (RHD), and the C-terminal rhodanese or sulfurtransferase domain (Rhod). Initial crystallographic structures of as-isolated CstB solved in the presence of reductant to 1.91 Å (Table S1) revealed the anticipated dimer-of-dimers tetrameric architecture; however, the C-terminal Rhod domain was not visible in these maps (Fig. S1a). The Fe(II) site was, however, well-defined by the data, revealing an octahedral Fe(II) coordination site, with three protein-derived ligands, H56, H119, and D145, arranged as a facial triad on one side of the Fe(II), and three open coordination sites modeled with high occupancy water molecules (Fig. S2; Table S2). There is no observable electron density for residues 197-207 in any subunit of the PDO domain, which contains C201 that previous studies implicate as playing an important role in catalysis (Fig. S1a)^19^. Finally, one of these structures reveals that C20, ≈29 Å away from each Fe(II) atom in the opposite protomer is persulfidated (PDB 28MT) or sulfonated (28MU), the significance of which is unknown.

In all remaining crystal structures of CstBs, solved in the absence of reductant, we were able to model both the C-terminal Rhod domain and the loop that harbors C201 in the PDO domain. WT Fe(II)-loaded CstB solved to 2.26 Å resolution (crystallographic statistics available in Table S1), adopts as expected the same dimer-of-dimers tetrameric architecture (Fig. 1a-c)^19^. The tetramer (Fig. 1a) is formed by interactions between the middle RHDs across the tetramer interface, with two PDO domains adopting a two-fold symmetric dimer at the “top” of each tetramer (see also Fig. S1b). The four C-terminal Rhod domains are positioned on the periphery of the tetramer and do not interact substantively with one another (Fig. 1a-b). The Fe(II) chelate, like in the first structure described (Fig. S2; Table S2), adopts a classical non-heme Fe(II) arrangement of ligands, involving a His_2_Asp triad of ligands. There is one axially positioned high-occupancy water molecule positioned at one of the open coordination sites that is observed in this wild-type CstB structure, hydrogen bonded to H58 (Fig. S2; Table S2). The entire CstB-specific loop is visible in these maps, with the S^γ^ of C201 forming the apex of this loop positioned 3.4 Å away from the Fe(II) and is thus not directly coordinated to the Fe(II) (Fig. 1d-f). Analysis of the crystallographic B-factors suggest that this loop is significantly more dynamic than the immediately surrounding regions of the PDO domain besides a solvent-exposed peripheral helical domain (Fig. 1c).

**Figure 1.**
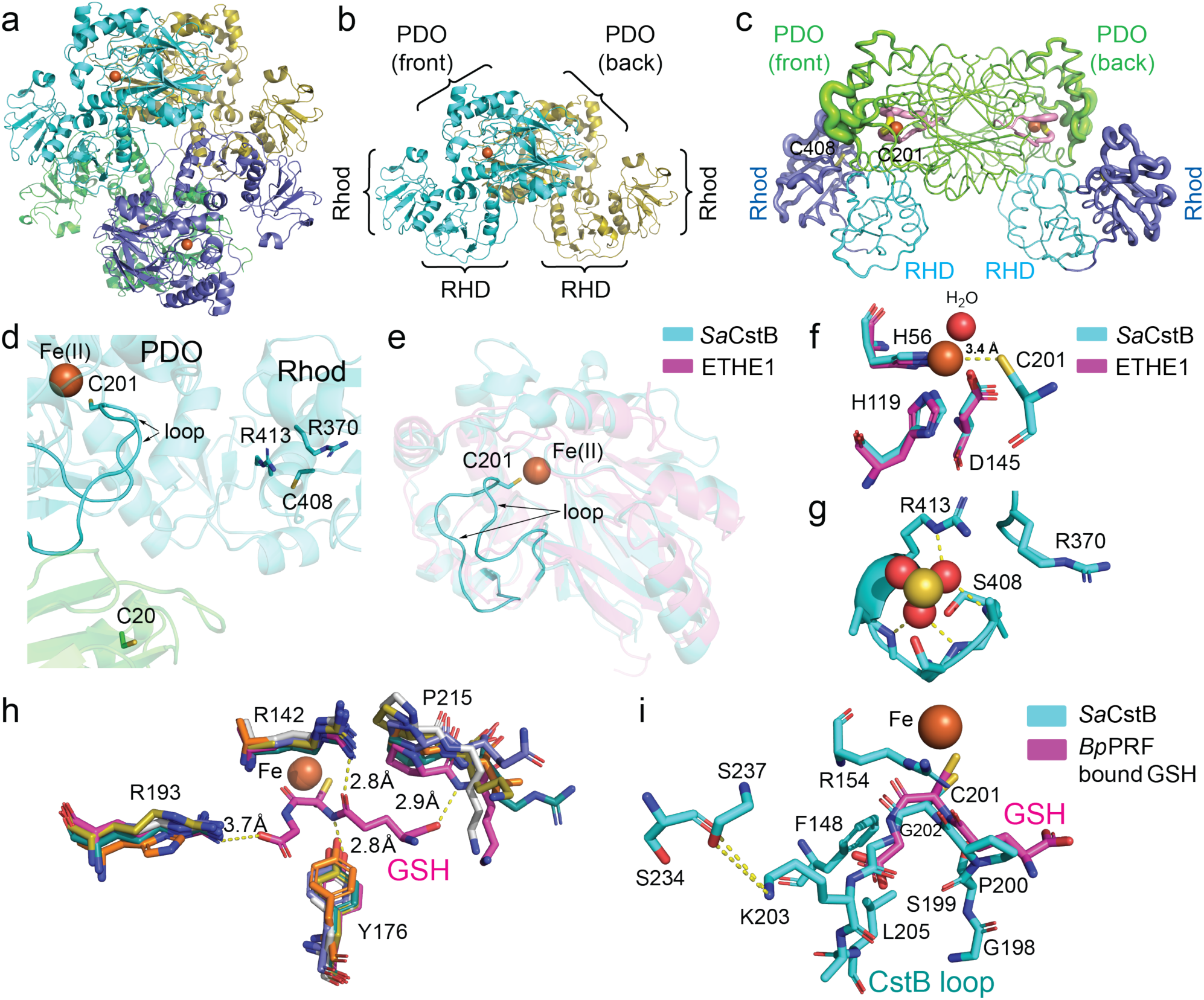
Structural characterization of CstB. **(a)** Ribbon representation of the crystallographic structure of the dimer-of-dimers *D*_2_ symmetric homotetramer of wild-type CstB (2.26 Å resolution) generated from the homodimer present in each unit cell (Table S1). Each subunit is shaded differently with the Fe(II) atoms in each PDO domain shaded *rust*. **(b)** Ribbon representation of the top AB dimer (*cyan-gold*) with the subdomains of each chain labeled. PDO, persulfide dioxygenase; RHD, rhodanese homology domain; Rhod, C-terminal rhodanese (sulfurtransferase) domain. **(c)** “Sausage-plot” illustration of the crystallographic Cα B-factors in the wild-type CstB homodimer, with the cartoon shaded according to domain, PDO (residues 1-256) (*green*), RHD (257-346) (*cyan*) and Rhod (347-444) (*blue*). The active site loop harboring C201 (yellow) is shaded pink, with C408 in the Rhod domain also highlighted (*yellow*). **(d)** Critical residues of CstB of interest in this study. The CstB-unique loop that harbors of the apical C201-C202 sequence is highlighted in *cyan* from the *cyan* shaded subunit with C20’ derived from the opposite dimer across the tetramer interface. **(e)** Structural alignment of CstB PDO domain with hETHE1, again highlighting the active-site loop in CstB that is not present in ETHE1. **(f)** Structural alignment of the ETHE1 (cyan) and CstB Fe(II) active sites (*magenta*), with the residue positions of CstB indicated. The side chain of C201 is also shown, unique to CstB. **(g)** Bound sulfite shown is spacefill in the Rhod domain of both C408S (shown) and C201S/C408S CstB mutants, with hydrogen bonding interactions shown in *yellow* dashed lines. **(h)** Superposition of the active site regions of all five structurally characterized GSSH-oxidizing PDOs with numbering based on the *Bp*PRF amino acid sequence (5VE5)^31^. Others are a plant ETHE1-like enzyme (2GCU)^34, 70^, human ETHE1 (4CHL)^33^, two bacterial enzymes (4YSB, 4YSK)^22^ and (8ZBD). The bound product GSH from the *Bp*PRF structure is shown in *magenta* as are selected intermolecular interactions specific to GSSH-oxidizing PDOs. **(i)** Wild-type CstB loop (PDB: 28MV) (*cyan*) and *Bp*PRF-bound GSH (PDB: 5VE5)^31^ (*magenta*) aligned with residue numbers based on the CstB amino acid sequence. Selected interactions between the active site loop and core of the PDO domain in CstB are shown.

To further investigate the close proximity of the S^γ^ of C201 to the Fe, Fe K-edge X-ray absorption spectroscopy (XAS) was employed to better understand the Fe coordination environment of C408A CstB (Fig. S3). The edge is characterized by a broad pre-edge with inflections at 7111.8 and 7114 eV, typical of octahedral Fe(II). An intense whiteline at ∼7124 eV is indicative of solely light atom (N/O) coordination. Extended X-ray Absorption Fine Structure (EXAFS) measurements were collected and analyzed to *k* = 12 Å^-1^. Analysis of the EXAFS region suggest a five- or six-coordinate Fe(II)-O/N complex, with one short N(His)/O scatterer ∼1.95 Å and four longer N/O scatterers at 2.12 Å. These features are consistent with the facial triad coordination environment found by crystallography with 2-3 additional H_2_O ligands (Table S3). Notably, a combination of XAS and EXAFS provide no evidence of direct coordination by S, consistent with the crystallographic 3.4 Å Fe-S_C201_ distance.

The interface formed between the Rhod and PDO domains is rather limited, largely involving residues 290-296 of the PDO domain and residues 415-425 of the Rhod domain within the same subunit (Fig. 1a-b). Crystallographic B-factors reveal that this interface is considerably dynamic, on par with much of the Rhod domain itself (Fig. 1c). This interdomain arrangement, however, creates an unobstructed space between S^γ^ atoms of C201 and C408, which are separated by 26.8 Å (Fig. 1d). It is interesting to note that both S^γ^ atoms are largely shielded from solvent. The N-terminal domain of CstB adopts the same fold as the human ETHE1 (Fig. S4a) despite a sequence identity of only 21 %^19^. The C-terminal rhodanese domain of CstB also displayed a similar fold with the human rhodanese (sulfurtransferase TSTD1) despite a meager 23 % sequence identity (Fig. S4a). C408 sits behind a positively charged “wall” formed largely by the side chains of R370 and R413; here, these two arginine residues create a patch of high positive charge density around this C408 S^γ^ atom (Fig. S5).

We also solved the crystallographic structures of the C408S, C201S/C408S, and C201S CstB mutants to 2.48 Å, 2.48 Å and 3.11 Å resolution, respectively, by molecular replacement with the wild-type enzyme (see Table S1 for crystallographic statistics). The overall fold of each of these structures is indistinguishable from the wild-type enzyme (Fig. S1b), but closer inspection reveals some key differences. The most notable difference is the presence of a sulfite ion bound to the rhodanese domain in both structures that harbor the C408S substitution (C408S; C201S/C408S), penetrating the positively charged “wall” around C408 (Fig. 1g; Table S2). The coordination chemistry around the Fe(II) in all three mutant structures is qualitatively similar to wild-type CstB, with some differences. The Fe(II) sites feature just one (C408S) or no (C201S/C408S; C201S) water molecules (Fig. S2; Table S2); and in the C201S/C408S structure, one unusually long metal-coordination bond, which differs in subunit A and subunit B. Otherwise, the metal-ligand coordination bond distances are wild-type like (2.1-2.2 Å). The C201-harboring loop is also well-ordered in all three structures with the same degree of dynamical disorder found in the wild-type enzyme. In both C408S-containing structures, the Fe(II)-C201 S^γ^ (or O^γ^) distance is elongated relative to wild-type CstB, to 3.6-3.7 Å, while in the C201S mutant, the S201 O^γ^ has rotated away from the Fe(II), adopting a distinct rotamer (5.2 Å away). Finally, there is additional electron density in the vicinity of C408, specifically in C201S CstB that is difficult to model with confidence but may represent a polysulfide chain on C408 (Fig. S6).

### CstB crystal structure predicts the role of the active-site loop

The human PDO, ETHE1, has a strong preference towards glutathione persulfide (GSSH) as the substrate and has no detectable or minimal activity against other persulfides such as cysteine persulfide (CSSH), homocysteine persulfide, and CoA persulfide (CoASSH)^20, 30^. The recently characterized persulfide dioxygenase–rhodanese fusion protein from *Burkholderia phytofirmans* (*Bp*PRF) also exhibits a similar preference for GSSH^31^, suggesting that the active sites of these PDOs have features that discriminate GSSH over other substrates. In fact, all PDOs crystallized thus far are GSSH-catalyzing^22, 31-34^, and alignment of their structures with bound GSH product reveals features shared by these enzymes that may stabilize the GSSH substrate in the active site (Fig. 1h). CstB lacks all conserved GSH-stabilizing features apart from R142 (R154 in CstB) (Fig. 1h), suggesting a disparate substrate preference for CstB. Remarkably, the C201-G202 dipeptide found in the long, flexible loop (consisting of residues 192 to 213) in CstB is nearly superimposable with the GSH product bound to *Bp*PRF, with the GSH and C201 cysteine sulfur atoms positioned in the same location relative to the Fe(II) site, in the axial coordination region (Fig. 1i). This observation hints at the plausible role of the C201-G202 dipeptide as a glutathione mimic, *i.e.*, a persulfidated C201 is the true substrate for this enzyme. The CstB loop is stabilized by several factors, including the H-bonding interaction of R154 to the P200 amide oxygen, the H-bonding interactions of K203 to the sidechains of S234 and S237 (Fig. 1i), as well as to N238 and S237-N238 amide backbone (not shown). Furthermore, F148 is stacked against L205, both of which are invariant in all CstB-like proteins (Fig. S7).

### CstB exhibits no substrate preference between GSSH and CSSH

The hypothesis that the C201-G202 dipeptide is a GSH mimic predicts that persulfide transfer between the LMW persulfide, *e.g*., CSSH, GSSH, to the C201-G202 dipeptide creates the substrate for Fe(II)-dependent oxidation. Furthermore, a low-molecular-weight (LMW) persulfide will not bind to CstB Fe active site in a saturable manner, making the identity of the LMW persulfide unimportant; *i.e.*, CstB has no substrate preference, unlike ETHE1 and *Bp*PRF^20, 31^. Consistent with this prediction, there is little discernible difference in the [RSSH]-dependence of the rate of O2-consumption between the two substrates (Fig. 2a). Furthermore, tetrasulfide (S_4_^2-^) also appears to be a viable substrate (Fig. S8), although this substrate substantially degrades when added to the reaction buffer. The observed activity with CSSH and GSSH appears linear over the persulfide concentration range of 0 to 100 µM and could not be fitted to a hyperbolic function as required by non-cooperative substrate binding in the Michaelis–Menten model. Beyond 100 µM substrate, the enzyme activity sharply declines with both substrates. We find that this inhibition is due to residual sulfide present in the thiol persulfide substrate cocktail (Fig. 2b)^20^. Indeed, there is no inhibition by the LMW thiol product even at 10 mM, consistent with the idea that thiols or persulfides do not bind to the Fe active site. Finally, the C201A enzyme is inactive in O2 consumption (Fig. 2c), consistent with a loss of “substrate” in this mutant, despite adopting a wild-type-like fold (Fig. S1b).

**Figure 2.**
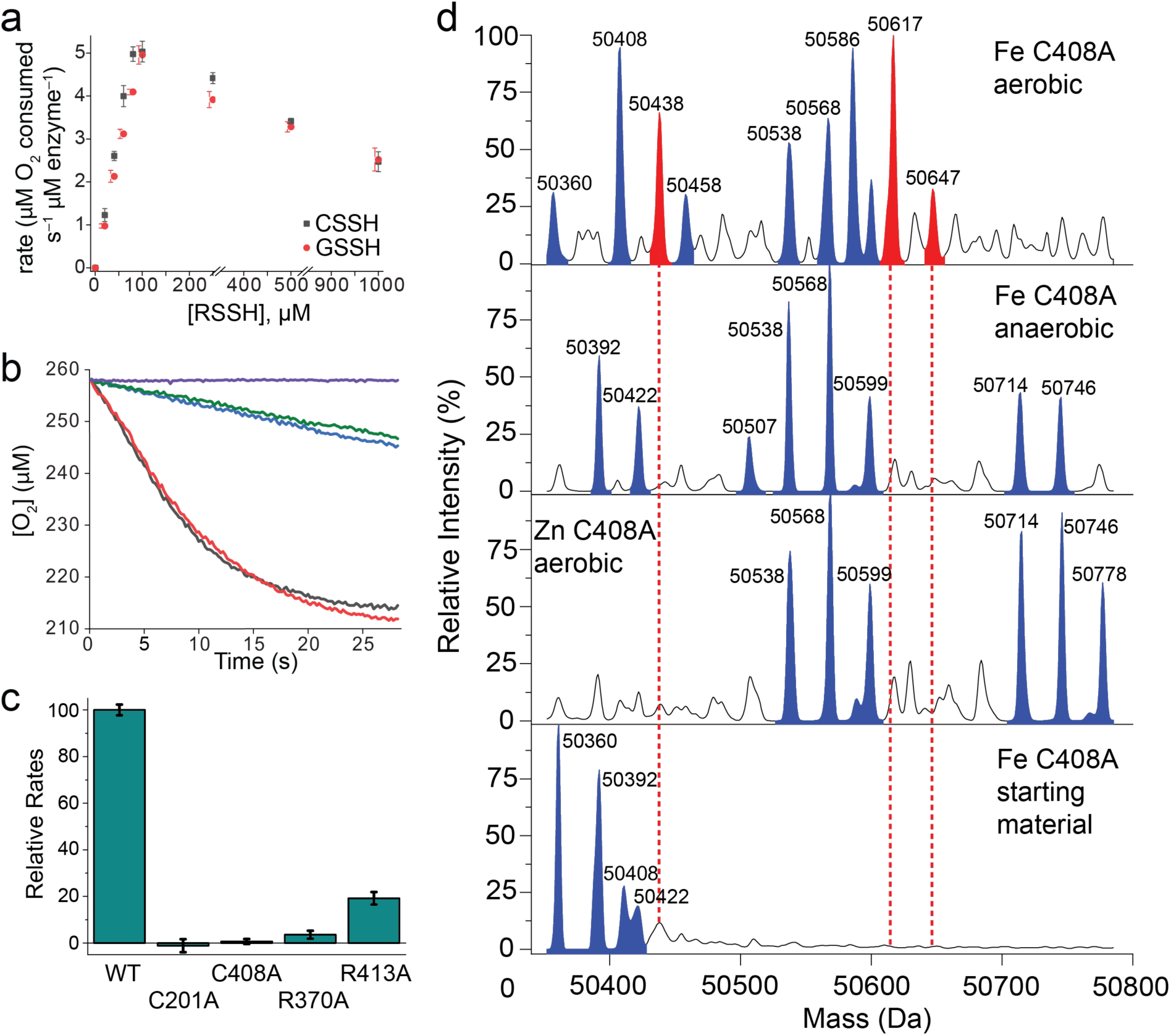
Enzyme turnover characteristics of CstB and variants. **(a)** Initial rate of oxygen consumption as a function of cysteine persulfide (CSSH) and glutathione persulfide (GSSH) concentration. **(b)** Representative rates of O_2_ consumption by wild-type CstB with no substate added (*violet trace*), with 100 µM CSSH (*black trace*), with 100 µM CSSH + 10 mM cysteine (*red trace*), with 100 µM CSSH + 10 mM sodium sulfide (*blue trace*), and with 10 mM sodium sulfide alone (*green trace*). **(c)** Bar chart of the relative initial rates of O_2_ consumption using 100 µM CSSH obtain for wild-type (WT) CstB and single-site variants, C201A, C408A, R370A, and R413A CstBs. **(d)** Deconvoluted intact ESI-MS mass spectra of C408A CstB loaded with Fe(II) or Zn(II) and incubated with CysSSH under aerobic or anaerobic conditions as indicated. Peaks without shading are considered background, while peaks shaded in *red* are indicative of masses consistent with an *S*-sulfonate modification. MS/MS spectra (Fig. S8) reveal that this *S*-sulfonate modification occurs specifically on a peptide containing C201. See Fig. S4 for a list of possible identifications of these mass species.

### C408A substitution abolishes the activity of the PDO domain

We previously reported that the C408S substitution negatively impacts the thiosulfate production of CstB and that no sulfite can be detected even when the PDO domain remains intact^19^. This finding suggests strong coupling between the PDO and the sulfurtransferase (Rhod) domain active sites. Our oxygen-consumption assay, which measures the activity of the PDO domain, confirms that the C408A variant like the C201S variant, is inactive, despite the CstB PDO domain remaining intact (Fig. 2c). Based on this result, we hypothesized that the sulfite intermediate in C408A remains tethered to the PDO domain as an *S*-sulfonate adduct to C201. To test this, we performed persulfidation of the Fe-loaded C408A variant, aerobically and anaerobically (Fig. 2d). We then capped any unreacted cysteine thiol and cysteine persulfide using β-(4-hydroxyphenyl)ethyl iodoacetamide (HPE-IAM) and analyzed the intact mass product distribution using ESI-MS. We detected cysteine persulfidation in both aerobic and anaerobic conditions, but it is more extensive for the anaerobic sample (Table S4; Fig. 2d). We also detected a species bearing what appears to be addition of a sulfonate (-SO_3_^-^ and -SSO_3_^-^) moiety, which is a significant product under aerobic conditions, but is not detected under anaerobic conditions (relevant masses: 50438 Da, 50617 Da, and 50586 Da; Table S4). This result is consistent with the hypothesis that this *S*-sulfonation reaction is strictly O_2_-dependent. We next performed the same experiment with the Zn-loaded C408A under aerobic conditions. The Zn-loaded aerobic enzyme exhibited similar mass spectral characteristics to the anaerobic Fe-loaded C408A, which confirms the observation of Fe-dependent oxidation. Finally, only the aerobic Zn-loaded and anaerobic Fe-loaded enzymes exhibit the addition of two HPE-IAM (relevant masses: 50714 Da, 50746 Da, and 50778 Da; Table S4), presumably because the two cysteines (C20 and C201) remain “cappable” by HPE-IAM only for these samples.

### A likely noncanonical two-electron reduced sulfonate mediates thiosulfate production

We next used MS/MS to unambiguously locate the various chemical modifications in the primary structure of C408A CstB upon incubation with the CSSH cocktail. Our analysis confirms that both C201 and C408 in an Fe-loaded sample prepared anaerobically can be persulfidated (Fig. S9a-b). More importantly, this analysis captures the existence of an *S*-sulfonated adduct covalently attached to C201 in the C408A variant only (Fig. S9c), revealing that the *S*-sulfonation is Fe and O_2_-dependent, a finding that is consistent with the parent masses (Fig. 2d). Interestingly, *S*-sulfonation on GSH has been previously synthesized as a PDO substrate analog, and it exhibits binding comparable to glutathione in ETHE1^22^, suggesting that *S*-sulfonation is a stable adduct. Persulfidation at C20 was also observed, in both Fe-loaded anaerobic and Zn-loaded aerobic samples, consistent with the fact that the two Cys (C20 and C201) can be persulfidated and capped, as revealed by the intact masses (Fig. 2d) and the structure (28MT) (Table S2).

We note that a canonical C201 *S*-sulfonation appears insufficient to produce thiosulfate upon reaction with a C408 persulfide (Fig. 3a), indicating that additional activation or structural modification of the sulfonate is required to form thiosulfate. Indeed, formation of an *S*-sulfonate from water and molecular oxygen, validated by product analysis (*vide infra*), is accompanied by the release of two electrons, as illustrated by the reaction:

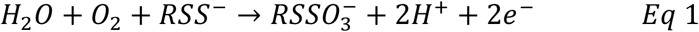

**Figure 3.**
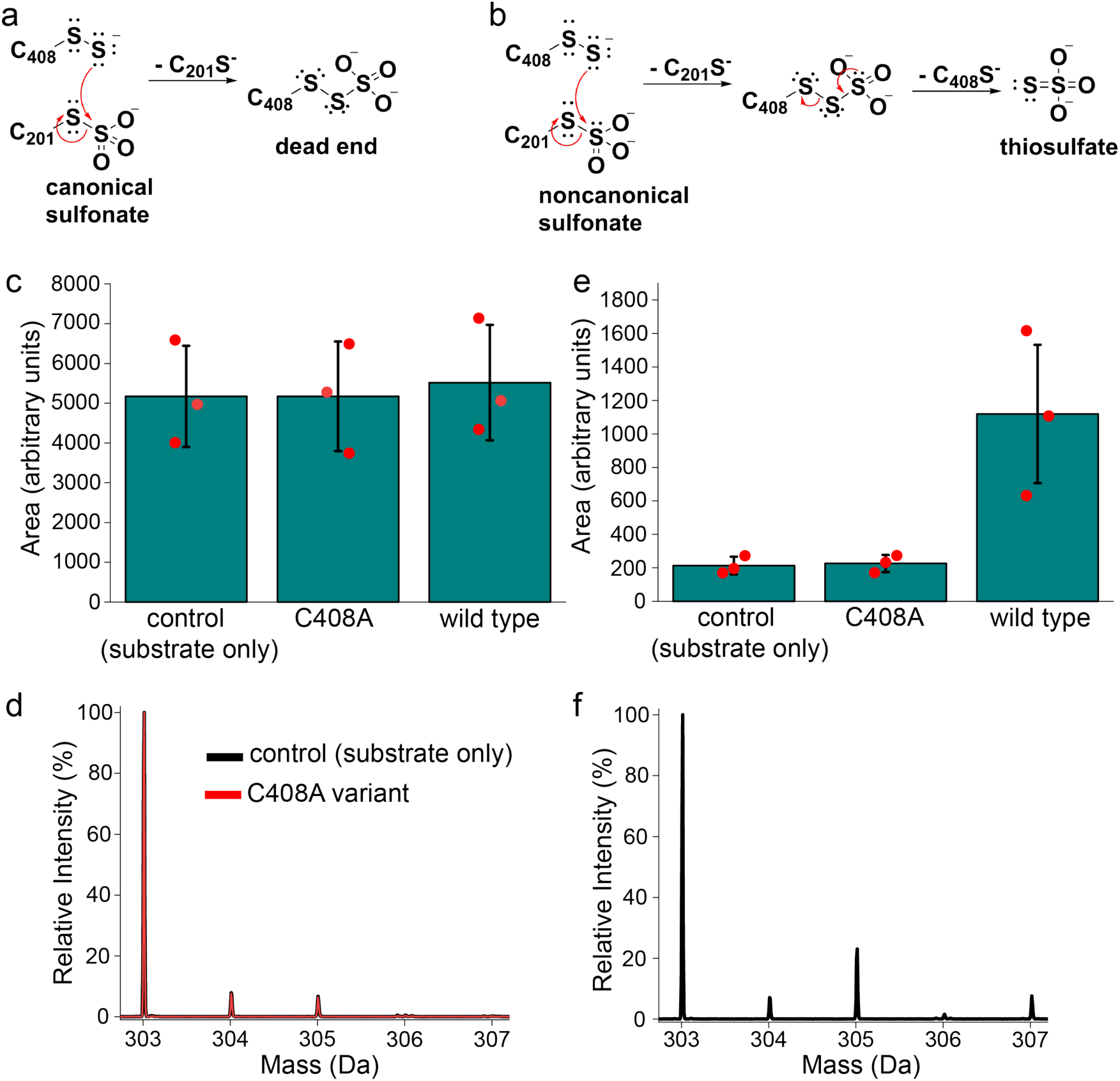
**(a)-(b).** Mechanistic scenarios illustrating the attack by the C408 persulfide on a **(a)** canonical or **(b)** two-electron reduced, non-canonical C201-*S*-sulfonate. Only the mechanism illustrated in panel (b) leads to the production of thiosulfate (TS) as product with regeneration of thiols at both C201 and C408. Other mechanistic scenarios are possible, including attack by contaminating sulfide in RSSH substrate cocktails on the S-S bond (Fig. S13). However, none are compatible with the stoichiometry of the enzyme-catalyzed reaction, where two mol of RSSH are consumed for every mol of TS made^19^. **(c)-(f).** Product analysis. Relative quantities of TS measured by integrating the extracted ion chromatograms (EICs) generated using the monoisotopic mass of **(c)** unlabeled and **(e)** ^18^O-labeled mBBr-thiosulfate adduct. Isotope distributions of the mBBr–thiosulfate adduct were also measured for **(d)** control (substrate only; no enzyme) and a reaction containing C408A CstB and **(f)** the analogous WT CstB-catalyzed reaction. The theoretical mass spectrum of “light” TS is shown in Fig. S10 for comparison.

Since the formation of Fe(0) from a two-electron reduction of Fe(II) is thermodynamically unfeasible, we hypothesize that the “released” electrons must be accommodated by the sulfonate group itself, forming a two-electron reduced noncanonical sulfonate, which is sufficient to enable thiosulfate formation after the nucleophilic attack by the C408 persulfide (Fig. 3b).

### Product analysis establishes the incorporation of one oxygen atom from water into the thiosulfate product

To validate the proposed reaction in Eq 1, which ultimately leads to the formation of TS (Fig. 3b), we performed product analysis using ^18^O-labeled water, and quantified the relative amounts of labeled and unlabeled sulfite and thiosulfate as monobromobimane (mBBr) adducts using ESI-MS. We could not detect a mBBr-sulfite adduct in assays with the wild type and C408A enzymes, consistent with the hypothesis that the catalytic mechanism does not involve the release of sulfite. We did detect the monoisotopic mass of unlabeled mBBr-thiosulfate adduct in the assay with the wild-type enzyme, but we observed similar levels of unlabeled mBBr-thiosulfate adduct with the C408A variant, and in the absence of enzyme (control, Fig. 3c). This observation suggests that the unlabeled thiosulfate is not a product of the enzymatic reaction but is a contaminant in the assay. Indeed, the normal isotope distribution of the mBBr-thiosulfate detected in the control and C408S samples (Fig. 3d) closely matches that of the theoretical isotope distribution of unlabeled TS (Fig. S10). In contrast, quantitation of the "heavy" ^18^O-labeled mBBr-TS adduct (exact monoisotopic mass: 305.015186 u) was considerably higher in reactions run with the wild-type enzyme, relative to the control and C408A variants (Fig. 3e); in addition, only the wild-type enzyme produced a deviation from the normal isotope distribution of “light” mBBr-TS adduct precisely at the monoisotopic mass corresponding to ^18^O-mBBr-TS adduct (Fig. 3f). These observations reveal that only the wild-type enzyme is active and that the oxidation of its C201 persulfide substrate involves the incorporation of an oxygen atom from water and two oxygen atoms from O_2_, to form ^18^O-labeled TS. Interestingly, a peak that is ∼2 Da heavier than the monoisotopic peak of ^18^O-labeled mBBr-thiosulfate adduct has an intensity that deviates from the normal isotope distribution (Fig. 3f, 307.0175 u). The origin of this is unknown but one possibility is that the Fe active site with a bound H_2_^18^O exhibits a kinetic isotope effect and prefers ^18^O=^16^O as the substrate. This was not investigated further.

### Electrostatic steering of the C201-*S*-sulfonate to the Rhod active site is important for activity

The failure of C408A CstB to turn over, coupled with the binding of sulfite to the Rhod domain of both C408S mutants, suggests the possibility that the strongly negatively charged *S*-sulfonated C201 is captured via electrostatic steering and subsequently processed by the C-terminal Rhod domain. This makes the prediction that substitution of major contributors to the positive electrostatic surface patch in the Rhod domain, R370 and R413, would negatively impact the rate of O_2_ consumption. This is what we find, as the R370A and R413A mutants are completely or significantly inactive, respectively (Fig. 2c). Furthermore, we find that the addition of sulfite to these reactions strongly inhibits the O_2_ consumption, presumably via competition for docking at the positive patch with *S*-sulfonated C201 (Fig. 1g; Fig. S11).

The Fe and Rhod active sites are separated by ≈27 Å within each subunit, and the kinetic data suggest that the *S*-sulfonated C201 migrates to the Rhod domain. It is interesting to note that B-factors for residues in this loop are elevated relative to much of the rest of the PDO domain (Fig. 1c). Furthermore, the loop harbors multiple glycine residues (G194, G196, G198, G202, and G206), with all but G202 in the glutathione mimic far from the active site Fe(II) (Fig. 4a). Of these five Gly residues, only G198, in all CstB structures, is characterized by a ϕ, ψ angle pair within the lower-right quadrant of the Ramachandran plot, where only Gly residues can be found due to steric considerations (Fig. 4b)^35^. The substitution of G198 with any amino acid is therefore predicted to restrict loop flexibility or otherwise alter what is likely an ensemble of loop conformations required for attack by the C408 persulfide. We find that the G198A mutant exhibits significantly lower O_2_ consumption activity at all [CSSH] tested, in striking contrast to the G206A mutant, which is only modestly affected (Fig. 4c). These data suggest that the G198A substitution negatively impacts oxygen chemistry at C201 and/or shuttling of the *S*-sulfonate to the Rhod domain.

**Figure 4.**
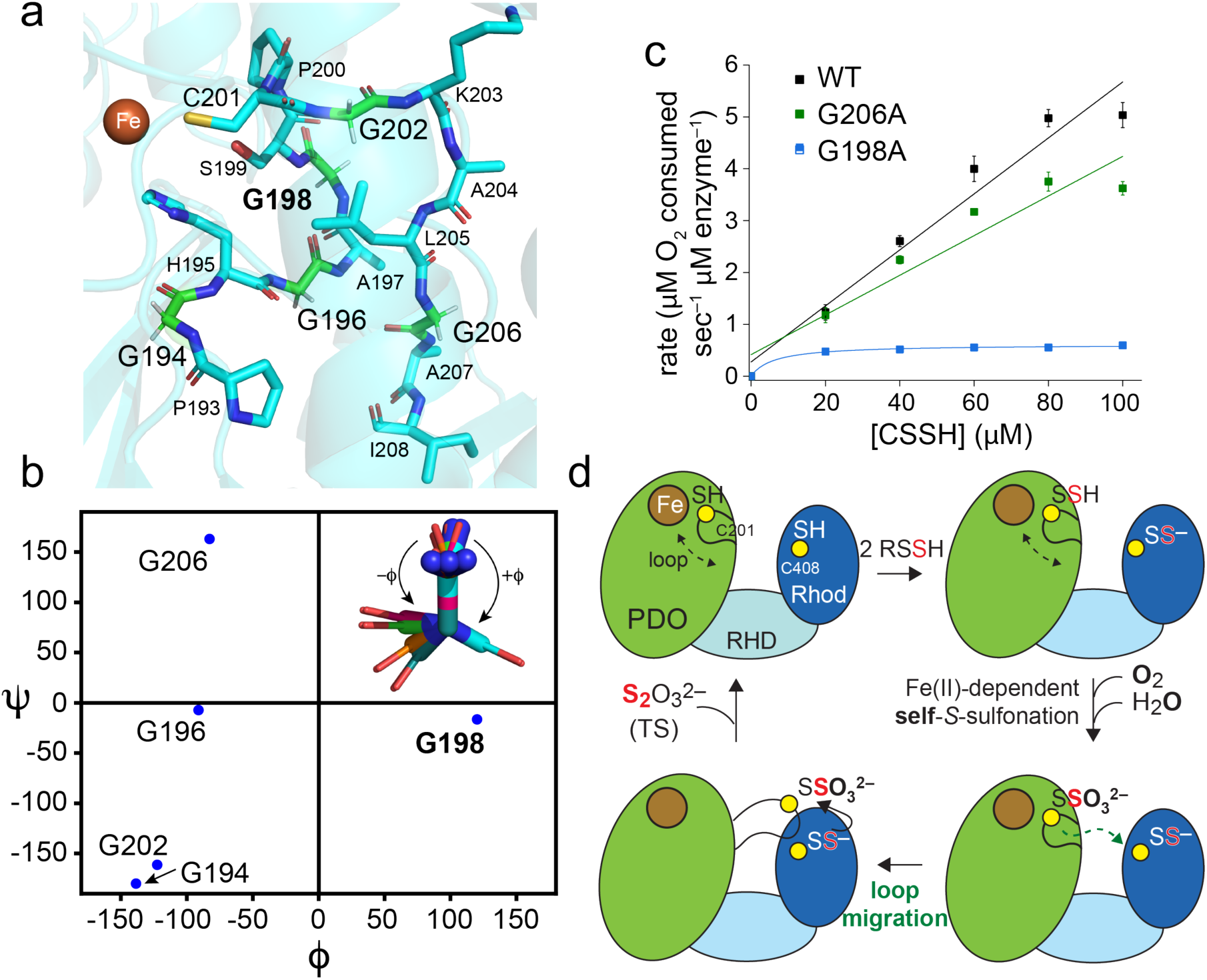
The active site loop migrates from the PDO to Rhod domain and couples O_2_ activation chemistry at C201 to the release of TS at C408. (**a**) The mobile loop (residues 193-208) of WT CstB contains five Gly residues, including the GSH mimic residue G202 and G198, which adopts phi (ϕ)-psi (ψ) angle space that can only be occupied by a Gly (panel b). (**b**) Ramachandran plot representation of the five Gly residues in the mobile loop. *Inset,* Illustration of the ϕ angle corresponding to all five Gly residues shown here; G198 (shaded *cyan*). **(c)** Rates of O_2_ consumption for the WT, G206A and G198A variant CstBs using CSSH as the substrate. (**d**) Cartoon model of how CstB processes LMW thiol persulfides (RSSH).

## Discussion

The persulfide dioxygenase, ETHE1, is strictly conserved in vertebrate mitochondria, highlighting the importance of sulfide homeostasis and prevention of the accumulation of harmful reactive sulfur species in eukaryotes. Given that glutathione is the most abundant thiol in the mitochondria^36, 37^, it is no surprise that ETHE1 exhibits high specificity for glutathione persulfide as its substrate. In contrast, *Staphylococcus aureus* contains five major low-molecular-weight thiols besides glutathione^38^, including bacillithiol, coenzyme A and cysteine^39^, and ergothioneine (data not shown), which together contribute to its cellular redox balance and defense against oxidative and other stressors, while maintaining CoASH acyl-transfer chemistry. *S. aureus* is known to colonize various sites of the human body, including the gastrointestinal tract known to have a high sulfide concentration. Therefore, *S. aureus* may have evolved a specialized mechanism that mitigates the toxicity of the various diverse persulfide species formed in a high sulfide environment of the gut, where in the presence of trace molecular oxygen, persulfide accumulation may reach levels that are toxic to cells. We hypothesized herein that *S. aureus* resolves this dilemma by employing CstB, a persulfide dioxygenase-rhodanese fusion (PRF), that we show is promiscuous towards persulfide substrates. This enzyme achieves this promiscuity primarily through rapid transpersulfidation of C201 in a CstB-unique loop relative to a canonical PDO, followed by molecular “shuttling” of a covalently bound sulfonate group to a persulfidated C408, and thiosulfate release. Indeed, the CstB Fe active-site lacks structural or electrostatic features that would stabilize the binding of LMW thiols and persulfides, such as those found in cysteine dioxygenase for binding its cysteine substrate^34^ or in canonical PDOs for binding the glutathione persulfide substrate.

When the persulfide attached to C201 in CstB is oxidized in an oxygen and Fe-dependent manner, it produces an *S*-sulfonate intermediate that remains bound to C201, instead of releasing free sulfite as is observed in a canonical PDO. This difference in the reaction product occurs even though the immediate Fe(II) coordination environment is the same in both enzyme types. The reason for this divergent outcome may lie in a specific hydrogen-bonding network, Thr---Asp---H_2_O, that is present in the canonical PDO active site but missing in CstB (Fig. S12a-b). This H-bonding network in canonical PDOs prevents the active-site water molecule from participating in the oxidation of the persulfide product, leading only to the formation of an RSSO_2_^-^ intermediate, which is then hydrolyzed by an external water molecule to release sulfite^40^. We considered the possibility that the Q→T substitution in CstB might abolish the H-bonding network thus unlocking an active-site water molecule hydrogen bonded to H58 (Fig. 1f; Fig. S2) that we propose facilitates the formation of the noncanonical *S*-sulfonate via a radical-based mechanism similar to that proposed for cysteine dioxygenase^41^ (Fig. 5). Efforts to switch CstB to a sulfite-producing ETHE1-like enzyme with a Q13T substitution failed since this enzyme is inactive (Fig. S12c); this suggests that other features must be required to switch the product distribution from TS to sulfite. We hypothesize that CstB evolved this mechanism to prevent the release of toxic sulfite, which interferes with the homeostasis of low molecular weight thiol homeostasis, impairing cellular antioxidant defenses^42^.

**Figure 5.**
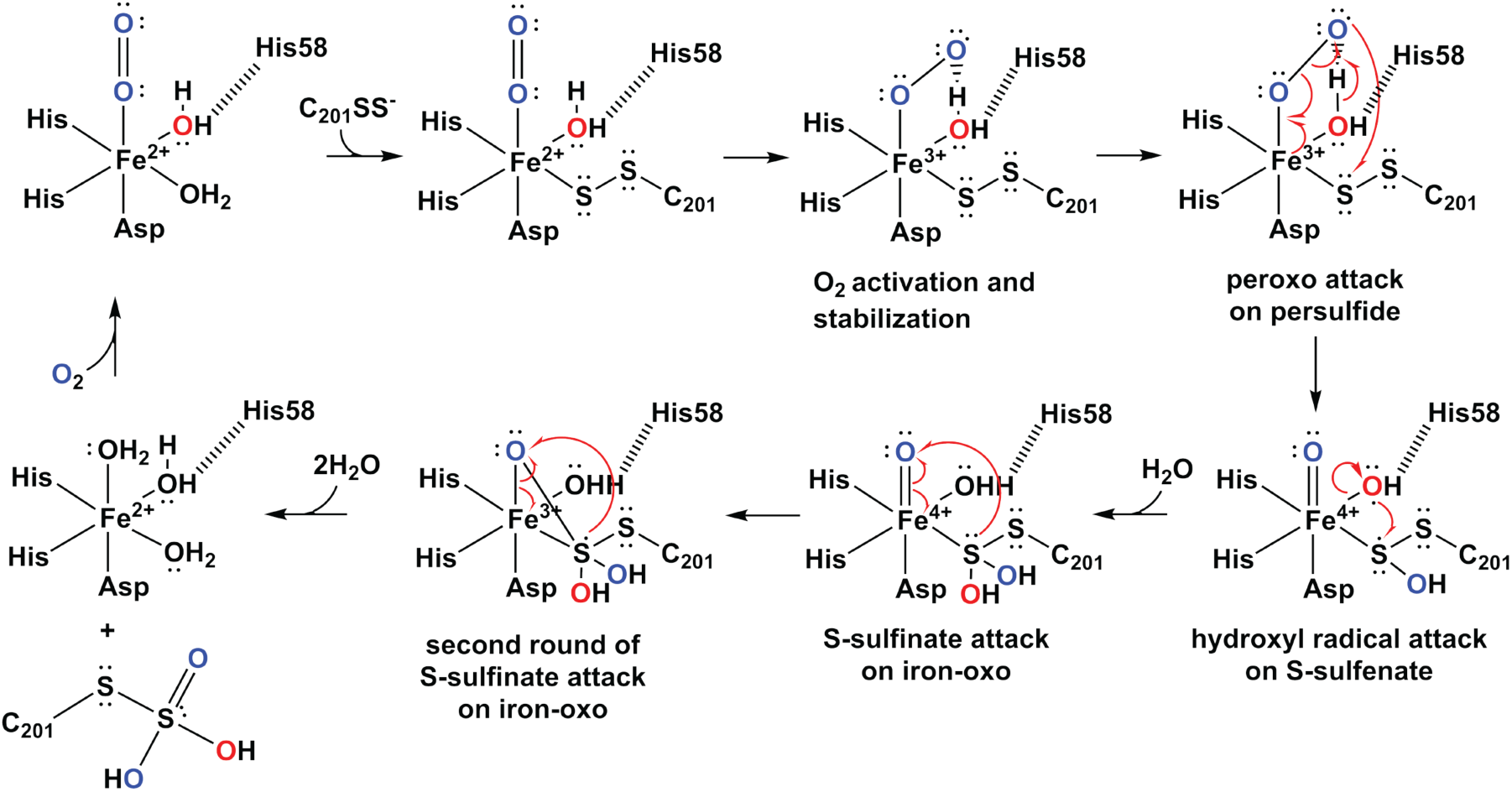
Plausible reaction mechanism that describes the installation of a noncanonical *S*-sulfonate at C201 at the Fe(II) active site in CstB that is consistent with the work presented here.

The *S*-sulfonate intermediate we uncovered in CstB is unlike the canonical sulfonate in several biological systems generated exclusively via sulfuransferases^43^. This sulfonate must be a two-electron reduced species, and although other mechanisms are possible (Fig. S13), each fails to recapitulate the 2:1 RSSH:TS stoichiometry observed in prior work^19^. Indeed, it is only in this two-electron reduced form that a reaction by a persulfide becomes productive, resulting in the formation of the TS product. We anticipate that the Rhod C408 persulfide (not the persulfide substrate itself) is the sole persulfide that can interact with the C201 *S*-sulfonate. This assumption is supported by the fact that the C408A variant is inactive and that the *S*-sulfonate formed under aerobic conditions remains tethered to C201 in this variant. Furthermore, the strong negative charge on the noncanonical *S*-sulfonate, as well as the presence of arginine residues R370 and R413 in the Rhod active site, suggests that the interaction between the PDO and Rhod domain is facilitated by electrostatic steering of the loop. Notably, we observed that the C408S variant crystallized with a bound sulfite in the Rhod active site even in the absence of sulfite added to the crystallization buffer. This suggests that the arginine residues surrounding the rhodanese active site play a crucial role in attracting and stabilizing a negatively charged moiety such as the noncanonical *S*-sulfonate. Interestingly, the *Bp*PRF C314S variant also crystallized with a chloride ion bound at its Rhod active site, where it is surrounded by arginine residues^31^, similar to the configuration observed in the CstB Rhod domain.

There is significant precedence for such a loop-mediated catalytic mechanism that we observe here in CstB. The iron-sulfur complex assembly (ISC) complex consists of the cysteine desulfurase sub-complex NFS1-ISD11-ACP1, the scaffold protein ISCU2, the electron donor ferredoxin FDX2, and frataxin, a protein dysfunctional in Friedreich’s ataxia^29^. The loop involved in the persulfide sulfur transfer from NFS1 to ISCU plays a crucial role as a dynamic "swinging arm" that facilitates the direct transfer of the sulfane sulfur to its acceptor. During *de novo* [2Fe-2S] cluster assembly, NFS1 generates a persulfide on a conserved cysteine residue within its flexible loop. This loop swings the persulfide from the NFS1 active site to the scaffold protein, ISCU2, where it is transferred onto a specific cysteine on ISCU2. The persulfidated ISCU2 then serves as the platform for Fe-S cluster assembly. Indeed, persulfides are generally thought to readily “shuttle” via a series of transpersulfidation reactions^27, 44^. A similar loop-mediated mechanism was also observed in CoAPR, where the pantothenate arm of a tightly bound coenzyme A molecule swings between two active sites of the enzyme^17^. The swinging arm brings the persulfide attached to a cysteine (C508) back to the reductase active site, where a second cysteine (C42) attacks it, releasing hydrogen sulfide (H_2_S) and forming a mixed disulfide intermediate. This intermediate is then reduced by flavin, which itself is reduced by NAD(P)H. The swinging arm allows for efficient substrate channeling and catalytic turnover preventing the release of intermediates. Such a “swinging arm” is rare for an unstructured element in enzymatic catalysis, and the “molecular shuttle” in CstB appears to be more functionally complex as it integrates oxidative and persulfide transferase activities in a highly concerted fashion.

## Materials and Methods

### Protein Expression and Purification

The full-length CstB from *Staphylococcus aureus* subsp. *aureus* strain Newman (locus tag: NWMN_0028) was codon-optimized, synthesized by Genscript, and inserted into the pET-28a(+) vector. The synthesized gene encodes wild-type CstB with a Strep-tag II sequence (WSHPQFEK)^45^ at the C-terminus. The CstB variants were then generated from this construct via site-directed mutagenesis, and the resulting plasmids were transformed into *E. coli* BL21(DE3) strain Rosetta. To overexpress the wild-type enzyme and its variants, 10 mL of an overnight culture (16-18 hours) was inoculated into 1 L of Luria-Bertani broth supplemented with 50 mg/mL kanamycin and 34 mg/mL chloramphenicol. The resulting culture was then incubated at 37 °C until the OD reached 0.6, after which the temperature was dropped to 18 °C and the culture was induced by adding IPTG to a final concentration of 1 mM. After 16–18 hours of incubation, cells were harvested by centrifugation and resuspended in freshly prepared lysis buffer (25 mM HEPES, 500 mM NaCl, 20 mM cysteine, pH 7.4). The resuspended cells were stored at -80 °C for at least 24 hours before protein purification.

To purify the enzyme, the frozen stock was warmed to room temperature to lyse the cells (*E.coli* BL21(DE3) strain Rosetta expresses lysozyme that facilitates cell lysis upon thawing after a freeze-thaw cycle). Before complete lysis, phenylmethylsulfonyl fluoride (PMSF) was added at a concentration of 5 mM along with a tablet of Protease Inhibitor Cocktail, mini-Tablet (EDTA-Free, MedChemExpress). The lysed cell suspension was then centrifuged at 208,000 rcf for 45 minutes, and the supernatant was separated and filtered using a 0.45-µm filter. The supernatant was loaded onto a Strep-Tactin XT 4flow high-capacity resin (IBA Lifesciences), after which the resin was washed with 15-20 ml of freshly prepared lysis buffer. The bound enzyme was eluted with 10 ml of 50 mM biotin dissolved in freshly made lysis buffer and then concentrated to approximately 2-3 ml. The concentrated enzyme was loaded onto a Superdex 200 Prep Grade column equilibrated with 25 mM HEPES, 150 mM NaCl, and 20 mM cysteine at pH 7.4. Finally, the purified enzyme was concentrated to approximately 200 µM, aliquoted into 20 μl volumes in PCR tubes, flash-frozen in liquid nitrogen, and stored at -80 °C.

### Protein Crystallization

Roughly, 500 µL of 9.5 mg/ml WT CstB and variants were dialyzed overnight in 25 mM MES at pH 6.0, containing 50 mM NaCl and 5 % (v/v) glycerol. The samples were further purified by size exclusion chromatography using a Superdex 200 Increase 10/300 GL (GE Healthcare Life Sciences) column with the same buffer. Our previous crystallization screening identified a buffer containing 20 % (V/V) PEG 3350, 0.1 M Bis-Tris propane pH 6.1-6.7, and 0.2 M sodium malonate (or sodium citrate) dibasic monohydrate as an optimal buffer for CstB. This time, the samples were screened using the same buffer at various PEG 3350 concentrations (17-23%), following the vapor diffusion sitting drop method^46^ at 20 °C and roughly 70 % humidity. Diffraction data for the wild type and C201S variant were collected at the ESRF (Grenoble, France), on ID23-2 and ID30A-3 (*MASSIF3*) beamlines with fixed wavelengths of 0.873 Å (wild type) and 0.968 Å (C201S variant). Diffraction data for the C408S and C201S/C408S variants were collected at the BESSY II Lightsource on BL14.1 beamline with a fixed wavelength of 0.918 Å.

Diffraction data were indexed and integrated using *XDS*,^47^ space-group assignment with *POINTLESS*,^48^ and scaling with *AIMLESS*^49^ and *STARANISO*^50^; all programs were used within the *autoPROC* data-processing pipeline.^51, 52^ Final diffraction data were converted to MTZ format with *CTRUNCATE*,^50, 53-55^ and a set corresponding to 5% of the total measured reflections was created and identified with Free-R flags.^56, 57^ Data quality was assessed with *phenix.xtriage* tool^58^ within the *PHENIX* suite of programs,^52, 59^ while the phasing of experimental data for wild-type *CstB* was done by molecular replacement using *PHASER*^60^ as implemented in *PHENIX*.^52, 59^ Two distinct search models were used: first, the previously determined “CstB incomplete model” (unpublished) for phasing the persulfide dioxygenase (PDO) and pseudorhodanese homology (RHD) domains; second, a rhodanese (Rhod) domain model generated using the I-TASSER server,^61^ constructed as a consensus model from 20 rhodanese structures deposited in PDB. Following the Matthews Coefficient analysis within *phenix.xtriage*, two molecules of each search model were used for phasing the experimental data (asymmetric unit defined as a dimer of the full-length *CstB*).

Iterative model building and refinement were carried out in a cyclic manner using *phenix.refine*^62^ within the *PHENIX* suite, *BUSTER-TNT*,^63^ and *COOT*^64^ until a complete model was built and refinement convergence achieved. The CstB models were validated using *MolProbity*^65^ as implemented in *PHENIX*. After initial molecular replacement phasing, phases for all subsequent *CstB* isomorphic datasets were determined through an initial rigid-body refinement with *BUSTER-TNT* using the corresponding apo dataset of each *CstB* variant. The “Missing Atoms” macro, together with the “-L” flag (“presence of an unknown ligand”), was used in *BUSTER-TNT* to search for unmodelled electron density on the maps. The structures were deposited in the Protein Data Bank under accession codes 28MT (truncated model, C20 persulfidated), 28MU (truncated model, C20 sulfonated), 28MV (full length model, “as isolated”), 28MW (full length, C201S variant), 28MX (full length, C408S variant), and 28MY (full length, C201S/C408S variant).

### Oxygen Consumption Assay

Anaerobic buffers for this experiment were degassed on a Schlenk line by purging 50 mL of buffer with ultrapure argon for 2 h then immediately transferred to a glove box for storage. The buffers contained 1 mM cysteine and were stored for at least four days before use. The added cysteine helps remove any residual O_2_ not eliminated by the degassing process. Aliquots containing sufficient enzyme for the experiment were thawed in the glove box and buffer-exchanged twice with buffer A containing 25 mM Tris, pH 8.0, 150 mM NaCl and 1 mM cysteine. The enzyme was diluted to approximately 500 µL (<100 µM concentration) and supplemented with 5 µL of 1 M FeSO_4_ to achieve a final Fe^2+^ concentration of 10 mM. For the Zn-loaded sample, 500 μL of 250 µM ZnSO_4_ was added to 100 µL protein. After incubating for 30 min, excess metal ions were removed by buffer exchanging three times with buffer A, followed by two exchanges with 100 mM sodium phosphate buffer (pH 7.4) containing 1 mM cysteine. Sodium dithionite was then added to a final concentration of 10 mM to reduce protein-bound iron and cysteine residues. Following a further 30-min incubation, the sample was buffer-exchanged four times with the same sodium phosphate buffer. The enzyme concentration was subsequently determined by UV–Vis spectroscopy using an extinction coefficient of 51,340 M⁻¹·cm⁻¹, which was calculated using Expasy ProtParam (https://web.expasy.org/protparam).

The oxygen consumption assay was performed using an oxygraph (Hansatech Instruments), which monitors oxygen uptake or evolution in real time using an S1 Clark-type electrode. The reaction mixture consisted of oxygen-saturated 100 mM sodium phosphate buffer (pH 7.4), 1 μM enzyme based on protein concentration, and persulfide substrate at the indicated concentration. The persulfide substrate was added first to the buffer, which was stirred at 70 rpm at 25 °C, and was quickly followed by the addition of the enzyme. Oxygen consumption was recorded until the reading plateaued. After the assay, the enzyme metal content was quantified by inductively coupled plasma mass spectrometry (ICP-MS), and the enzymatic activity was corrected on a per metal basis. The persulfide substrates were generated as previously described, with minor modifications^20^. Briefly, anaerobic solutions of 250 mM sodium sulfide and 50 mM oxidized cysteine (or glutathione), both dissolved in 300 mM HEPES buffer (pH 7.4), were prepared and mixed inside a glove box for 30 min. The resulting solution was distributed into 50 μL aliquots, flash-frozen in liquid nitrogen, and stored at –80 °C until use. Persulfide substrate concentrations were determined by cyanolysis^66^ with yields of roughly 50%.

### C408A CstB Persulfidation

The C408A enzyme was metalated with either iron or zinc as described above. The persulfidation reaction was performed in 100 mM sodium phosphate buffer (pH 7.4) containing 10 μM enzyme and 300 μM cysteine persulfide. For Fe-loaded C408A, the reaction was incubated for 10 min under both aerobic and anaerobic conditions. For Zn-loaded C408A, incubation was done only under aerobic conditions. After the 10-min incubation, β-(4-hydroxyphenyl)ethyl iodoacetamide (HPE-IAM) was added to 20 mM, and the mixtures were incubated for 1 h.

### Mass Spectrometry

The persulfidated samples described above were immediately analyzed by liquid chromatography coupled to electrospray ionization mass spectrometry on a Waters Synapt G2S mass spectrometer equipped with an iClass Acquity HPLC. Proteins were bound via a C4 reverse-phase column and eluted to determine the intact mass and modification state of the enzyme. To identify the site of modification in the enzyme, MS/MS analysis was performed. The samples were digested with trypsin at a 1:20 mass ratio and incubated at 37 °C for no more than 4 h. The aerobic Fe-loaded C408A sample prepared for LC-MS/MS analysis was not treated with β-(4-hydroxyphenyl)ethyl iodoacetamide (HPE-IAM) because the *S*-sulfonate adduct can only be detected in the absence of HPE-IAM and when trypsin digestion is limited to 2 h. The LC-MS/MS analysis was performed using a Orbitrap Fusion Lumos Tribrid mass spectrometer in the same manner as before^67^ except that the mass spectrometer was operated at a mass range between 350 to 2000 *m/z*, and the precursor ions were selected for MS/MS analysis at a 32% collision energy with an intensity threshold at 4.0 x10^5^. In addition, the dynamic exclusion duration was set at 30 s. Finally, mass tolerances for precursor and fragment ions were set to 10 ppm and 0.05 Da, respectively.

### CstB Product Analysis

The product analysis was performed similarly to the oxygen-consumption assay, except that the reaction was carried out in approximately 80% ^18^O-labeled water and without using an oxygraph. Each 20-μL reaction mixture contained 1 μM enzyme and 100 mM cysteine persulfide substrate. The reaction was allowed to proceed for 30 s and was terminated by adding 80 μL of a methanolic quenching solution, which consisted of 375 μL of 100 mM Tris buffer (pH 9.5) with 0.2 mM EDTA and 625 μL methanol. The resulting mixture was then added with 6 μL of 184.5 mM monobromobimane (mBBr) and incubated for 1 h at room temperature in a dark room. The reactions were either frozen at –80 °C for later analysis or immediately centrifuged at 4 °C using an ultracentrifugal filter with a 3 kDa molecular weight cut-off. This process filters out the protein component, allowing recovery of the filtrate for further analysis. The relative amounts of mBBr-sulfite and mBBr-thiosulfate adducts in the filtrate, both unlabeled and ^18^O-labeled, were analyzed by LC-MS using Waters Synapt G2S mass spectrometer equipped with a YMC-Triart C18 column. From the total ion chromatogram (TIC), extracted ion chromatograms (EICs) for unlabeled and ^18^O-labeled mBBr-sulfite and mBBr-thiosulfate were generated by selecting the exact monoisotopic *m/z* values of the target ions within a mass tolerance of 10 ppm. For each EIC generated, the area under each peak was integrated to quantify the relative amount of the target ion.

### **X-** ray Absorption Spectroscopy

Fe K-edge XAS measurements were acquired at Brookhaven National Laboratory (BNL) National Synchrotron Light Source II (NSLS-II) 8-ID ISS beamline^68^. A liquid N_2_-cooled double-crystal monochromator with Si(111) crystals was used to select the incoming X-ray energy with an intrinsic resolution (ΔE/E) of ∼0.6 x 10^-4^. The X-ray beam size was 1 mm x 1 mm (v x h) at the sample position. Samples were maintained at cryogenic temperatures using a closed cycle liquid He coldhead cryostat (∼15 K) to minimize radiation damage and maintain an inert sample environment. Partial fluorescence yield measurements were recorded using a 4-element Si Drift Diode (SDD) detector. Prior to measurements, each sample was checked for signs of radiation damage by performing subsequent 20 min scans over the same sample spot. All measured samples showed very low sensitivity to photoreduction. Scans were calibrated by simultaneous measurement of an Fe foil, with the first inflection point set to 7111.2 eV.

Individual XAS scans were normalized to the incident photon flux, energy calibrated, and merged using local processing software available at beamline 8-ID. Background subtraction and normalization were performed using a linear regression for the pre-edge region from 7020.6 to 7090.0 eV, and a quadratic polynomial regression for the post-edge region from 7270.0 to 7988.8 eV. Fe K-edge XAS spectra were splined from *k* = 0–15 Å^-1^ using an R-background of 1.0 and *k*-weight of 2. EXAFS fitting was performed using FEFF6 via the software package Demeter^69^. Possible scattering paths for EXAFS models were initially determined using FEFF6 in combination with the determined crystal structure. For all fits, the parameters R (bond distance), σ^2^ (bond variance), and *E_0_* (ionization energy) were allowed to vary during fitting refinement. A fixed value of S_0_^2^ = 1.0 was used in all EXAFS fits. A *k*-range of 3-12 Å^-1^ and R-range of 1-3 Å was used in the curve fitting analysis of all spectra, providing a maximum resolution of ΔR = 0.174 Å (according to the Rayleigh criterion) and 11.46 degrees of freedom based on the Nyquist criterion.

## Acknowledgements

We thank members of the Giedroc laboratory past and present for many helpful discussions over the course of this work. This work was supported by a grant from the US National Institutes of Health (R35 GM118157) to D.P.G. B.J.C.W was supported by the Training Program in Quantitative and Chemical Biology (QCB) at Indiana University (T32 GM131994).

## Author contributions

D.P.G., J.O.C. and J.A.B. designed the project. J.O.C., S.S.S, B.J.C.W., A.T.P. and J.K. performed all the experiments. J.O.C., J.C.T, C.V.S. and J.A.B. carried out the data analysis and interpretation, with overall supervision provided by D.P.G., G.G.-G. and M.A. The manuscript was written by J.O.C. and D.P.G. Funding was obtained by D.P.G.

