## Supporting Information for "Self-*S*-sulfonation in a bacterial persulfide dioxygenase mediates thiol persulfide detoxification"

This pdf file contains **Tables S1-S4**, **Figs. S1-S13** and Supplementary References.

**Table S1.** Data collection, processing, refinement statistics, and model quality parameters for *S. aureus* wild-type CstB (28MT, 28MU and 28MV), and the C201S (28MW), C408S (28MX), and C201S/C408S (28MY) CstB variants.

| | **Truncated model: C20 persulfurated** | | **Truncated model: C20 sulfonated** | | **Full length model: "as isolated"** | | **Full length model: C201S mutant** | | **Full length model: C408S mutant** | | **Full length model: C201S/C408S mutant** | |
|---|---|---|---|---|---|---|---|---|---|---|---|---|
| **PDB entry** | 28MT | | 28MU | | 28MV | | 28MW | | 28MX | | 28MY | |
| **Data collection** | | | | | | | | | | | | |
| **Synchrotron** | ESRF (Grenoble – France) | | ESRF (Grenoble – France) | | ESRF (Grenoble – France) | | ESRF (Grenoble – France) | | BESSY (Berlin – Germany) | | BESSY (Berlin – Germany) | |
| **Beamline** | ID30B | | ID30B | | ID23-2 | | ID30A-3 (*MASSIF3*) | | BL14.1 | | BL14.1 | |
| **Wavelength (Å)** | 1.000 | | 1.000 | | 0.873 | | 0.9677 | | 0.9184 | | 0.9184 | |
| **Space group** | $P\ 4_3\ 2_1\ 2$ | | $P\ 4_3\ 2_1\ 2$ | | $P\ 4_3\ 2_1\ 2$ | | $P\ 4_3\ 2_1\ 2$ | | $P\ 4_3\ 2_1\ 2$ | | $P\ 4_3\ 2_1\ 2$ | |
| **Unit cell** | | | | | | | | | | | | |
| ***a, b, c*** **(Å)** | 114.78, 114.78, 132.96 | | 115.28, 115.28, 133.20 | | 148.83, 148.83, 124.70 | | 147.09, 147.09, 125.09 | | 147.29, 147.29, 124.22 | | 147.65, 147.65, 124.90 | |
| ***α, β, γ*** **(°)** | 90, 90, 90 | | 90, 90, 90 | | 90, 90, 90 | | 90, 90, 90 | | 90, 90, 90 | | 90, 90, 90 | |
| | ***AIMLESS*** | ***STARANISO*** | ***AIMLESS*** | ***STARANISO*** | ***AIMLESS*** | ***STARANISO*** | ***AIMLESS*** | ***STARANISO*** | ***AIMLESS*** | ***STARANISO*** | ***AIMLESS*** | ***STARANISO*** |
| **Resolution range (Å)**[a] | | 86.89 – 1.95 (2.27 – 1.95) | 19.56 – 1.97 (2.01 – 1.97) | 19.77 – 1.91 (2.09 – 1.91) | 19.94 – 2.59 (2.63 – 2.59) | 19.42 – 2.26 (2.58 – 2.26) | 95.29 – 3.27 (3.33 – 3.27) | 95.29 – 3.11 (3.29 – 3.11) | 19.83 – 2.54 (2.58 – 2.54) | 19.93 – 2.48 (2.66 – 2.48) | 104.40 – 2.85 (2.90 – 2.85) | 104.40 – 2.48 (3.01 – 2.48) |
| **Total no. of reflections** | | 208 794 (10 338) | 635 561 (31 714) | 564 744 (31 364) | 282 962 (14 393) | 298 193 (19 659) | 368 739 (9 401) | 396 001 (26 150) | 610 388 (30 754) | 534 451 (27 340) | 386 769 (14 393) | 331 429 (8 144) |
| **No. of unique reflections** | | 32 247 (1 612) | 63 504 (3 133) | 55 925 (2 796) | 41 288 (2 057) | 40 674 (2 034) | 21 794 (1 068) | 22 176 (1 109) | 45 590 (2 213) | 40 074 (2 004) | 30 644 (1 603) | 25 782 (1 289) |
| **Multiplicity** | | 6.5 (6.4) | 10.0 (10.1) | 10.1 (11.2) | 6.9 (7.0) | 7.3 (9.7) | 16.9 (8.8) | 17.9 (23.6) | 13.4 (13.9) | 13.3 (13.6) | 12.6 (9.0) | 12.9 (6.3) |
| **Completeness (spherical) (%)** | | 49.3 (6.9) | 99.9 (100.0) | 79.4 (17.0) | 93.4 (94.7) | 62.1 (9.7) | 99.9 (99.9) | 87.9 (28.8) | 100.0 (100.0) | 81.7 (21.8) | 93.6 (99.2) | 51.9 (5.9) |
| **Completeness (ellipsoidal) (%)** | - | 90.9 (58.8) | - | 95.7 (59.5) | - | 84.9 (56.1) | - | 94.7 (53.2) | - | 95.7 (63.3) | - | 85.8 (48.3) |
| **Mean I/sigma(I)** | | 8.8 (2.1) | 11.0 (0.8) | 12.5 (1.5) | 7.6 (1.2) | 7.7 (1.5) | 7.4 (1.1) | 7.6 (1.2) | 8.2 (0.9) | 9.2 (1.4) | 7.7 (0.7) | 9.1 (1.4) |
| **Wilson B-factor ($Å^2$)** | - | 28.71 | - | 31.91 | - | 42.23 | - | 75.79 | - | 41.75 | - | 45.36 |
| $R_{merge}$ **(%)**[b] | | 14.9 (93.3) | 11.4 (267.0) | 10.4 (150.4) | 21.2 (206.1) | 22.2 (203.4) | 61.5 (229.0) | 71.4 (744.1) | 31.5 (330.7) | 27.6 (208.2) | 29.7 (282.2) | 25.7 (132.8) |
| $R_{meas}$ **(%)**[c] | | 16.2 (101.4) | 12.0 (281.2) | 11.0 (157.5) | 22.7 (221.2) | 23.8 (213.8) | 63.2 (242.4) | 73.4 (760.0) | 32.8 (343.3) | 28.7 (216.3) | 31.0 (299.1) | 26.8 (143.8) |
| $R_{pim}$ **(%)**[d] | | 6.3 (39.2) | 3.8 (87.1) | 3.4 (46.6) | 7.8 (76.5) | 8.0 (64.3) | 14.2 (75.8) | 16.1 (152.1) | 8.9 (91.7) | 7.8 (58.3) | 8.6 (94.3) | 7.4 (52.9) |
| $CC_{1/2}$ **(%)**[e] | | 99.6 (68.9) | 99.9 (34.3) | 99.9 (63.4) | 98.9 (39.) | 99.0 (58.2) | 97.0 (41.1) | 97.0 (36.5) | 99.3 (33.9) | 99.4 (54.9) | 95.4 (45.1) | 91.9 (46.0) |

| | Truncated – C20 persulfurated | | Truncated – C20 sulfonated | | Full length – “as isolated” | | Full length – C201S mutant | | Full length – C408S mutant | | Full length – C201S/C408S mutant | |
|---|---|---|---|---|---|---|---|---|---|---|---|---|
| **PDB entry** | 28MT | | 28MU | | 28MV | | 28MW | | 28MX | | 28MY | |
| **Refinement** | | | | | | | | | | | | |
| | ***AIMLESS*** | ***STARANISO*** | ***AIMLESS*** | ***STARANISO*** | ***AIMLESS*** | ***STARANISO*** | ***AIMLESS*** | ***STARANISO*** | ***AIMLESS*** | ***STARANISO*** | ***AIMLESS*** | ***STARANISO*** |
| $R_{cryst}$ **(%)**[f] | - | 17.81 (37.36) | - | 17.07 (33.91) | - | 17.73 (28.57) | - | 17.74 (29.35) | - | 18.14 (28.66) | - | 19.68 (39.06) |
| $R_{free}$ **(%)**[g] | - | 22.44 (39.17) | - | 20.15 (49.57) | - | 20.38 (39.80) | - | 21.93 (35.18) | - | 21.16 (26.86) | - | 23.15 (50.72) |
| **Number of atoms** | - | 5 286 | - | 5 378 | - | 7 190 | - | 7 037 | - | 7 221 | - | 7 093 |
| **Protein** | - | 5 034 | - | 5 032 | - | 7 023 | - | 7 023 | - | 7 039 | - | 7 046 |
| **Ligands** | - | 3 | - | 19 | - | 7 | - | 12 | - | 28 | - | 17 |
| **Waters** | - | 249 | - | 327 | - | 160 | - | 2 | - | 154 | - | 30 |
| **RMSD bonds (Å)**[h] | - | 0.004 | - | 0.004 | - | 0.003 | - | 0.002 | - | 0.002 | - | 0.002 |
| **RMSD angles (°)** | - | 0.64 | - | 0.69 | - | 0.64 | - | 0.54 | - | 0.51 | - | 0.58 |
| **Protein residues** | - | (G1 – P135)<br>(T168 – G196)<br>(A207 – S237)<br>(P243 – P345) | - | (G1 – P135)<br>(T168 – G196)<br>(A207 – S237)<br>(P243 – P345) | - | (M0 - K444)<br>[N(-7) - L443] | - | (M0 - K444)<br>[N(-7) - L443] | - | (M0 - K444)<br>[T(-9) - L443] | - | (M0 - K444)<br>[T(-10) - L443] |
| **Ramachandran plot** | | | | | | | | | | | | |
| **Most favoured (%)** | - | 97.23 | - | 97.42 | - | 95.85 | - | 95.85 | - | 95.75 | - | 95.75 |
| **Allowed (%)** | - | 2.77 | - | 2.58 | - | 4.04 | - | 4.04 | - | 4.14 | - | 4.14 |
| **Outliers (%)** | - | 0.00 | - | 0.00 | - | 0.11 | - | 0.11 | - | 0.11 | - | 0.11 |
| ***B*-factors (Å²)** | - | 33.63 | - | 40.73 | - | 50.84 | - | 75.52 | - | 47.28 | - | 47.40 |
| **Protein** | - | 33.891 | - | 40.73 | - | 51.02 | - | 75.52 | - | 47.50 | - | 47.46 |
| **Ligands/ions** | - | 35.18 | - | 50.43 | - | 47.13 | - | 80.59 | - | 55.96 | - | 49.60 |
| **Waters** | - | 30.09 | - | 40.13 | - | 42.85 | - | 40.77 | - | 35.71 | - | 32.12 |

[a] Information in parenthesis refers to the last resolution shell

[b] $R_{merge} = \Sigma_h\Sigma_l\,|I_{hl} - \langle I_h\rangle|/\Sigma_h\Sigma_l\,\langle I_h\rangle$, where $I_{hl}$ is the $l$th observation of reflection h and $\langle I_h\rangle$

[c] $R_{meas} = \Sigma_{hkl}(1/N{-}1)^{1/2}\Sigma_i|I_i(hkl) - (\overline{I(hkl)}|/\Sigma_{hkl}\Sigma_i I_i(hkl)$

[d] $R_{p.i.m} = \Sigma_{hkl}(1/N{-}1)^{1/2}\Sigma_i|I_i(hkl) - \overline{(I(hkl)}|/\Sigma_{hkl}\Sigma_i I_i(hkl)$

[e] $CC_{1/2}$ as described.[1]

[f] $R_{cryst} = \Sigma_h||F_{obs(h)}| - |F_{cal(h)}||/\Sigma_h|F_{obs(h)}$, where $F_{obs(H)} - F_{cal(h)}$ are the observed and calculated structure factors for reflection $h$, respectively.

[g] $R_{free}$ was calculated the same way as $R_{factor}$ but using only 5% of the reflections which were selected randomly and omitted from refinement

[h] RMSD, root mean square deviation

**Table S2.** Notable structural characteristics of all CstB structures reported in this work[a]

| PDB code | Variant | Resolution | Fe coordination | Cys20 modification | Ligands bound near C408 | Other notes |
|---|---|---|---|---|---|---|
| 28MT | WT: Truncated model | 1.95 Å | Hemifacial, three or no waters **Chain A:** three waters; **Chain B:** no waters | persulfidated | none | Rhod domains not visible (res 1-345); C201 loop not visible (res 197-206); another loop not visible (res 156-167) |
| 28MU | WT: Truncated model | 1.91 Å | Hemifacial, two or three waters **Chain A:** H56, 2.2 Å; H119, 2.2 Å; D145, 2.1 Å; waters, 2.2, 2.2, 2.1 Å **Chain B:** H56, 2.2 Å; H119, 2.2 Å; D145, 2.1 Å; waters, 2.2, 2.1 Å | sulfonated | none | Rhod domains not visible (res 1-345), C201 loop not visible (res 197-206); another loop not visible (res 156-167) |
| 28MV | WT: Full length model | 2.26 Å | Hemifacial, one water **Chain A:** H56, 2.1 Å; H119, 2.1 Å; D145 2.0 Å, water, 2.2 Å **Chain B:** H56, 2.1 Å; H119, 2.2 Å; D145 2.2 Å, water, 2.2 Å | none | none | **Chain A:** C201 S–Fe, 3.4 Å; **Chain B:** C201 S–Fe, 3.4 Å |
| 28MX | C408S: Full length model | 2.48 Å | hemifacial, one water **Chain A:** H56, 2.2 Å; H119, 2.2 Å; D145 2.1 Å, water, 2.1 Å **Chain B:** H56, 2.2 Å; H119, 2.2 Å; D145 2.1 Å, water, 2.1 Å | none | $SO_3^{2-}$ [b] | **Chain A:** C201 S–Fe, 3.7 Å **Chain B:** C201 S–Fe, 3.6 Å |
| 28MY | C201S/C408S: Full length model | 2.48 Å | Distorted hemifacial, no waters **Chain A:** H56, 2.3 Å; H119, 2.4 Å; **D145 2.9 Å** **Chain B**: H56, 2.3 Å; **H119, 2.8 Å;** D145 2.3 Å | none | $SO_3^{2-}$ [b] | **Chain A:** S201 O–Fe, 3.6 Å; H58 N–Fe, 4.3 Å **Chain B:** S201 O–Fe, 3.7 Å; H58 N–Fe, 4.3 Å |
| 28MW | C201S: Full length model | 3.11 Å | Hemifacial, no waters **Chain A:** H56, 2.2 Å; H119, 2.2 Å; D145, 2.0 Å **Chain B:** H56, 2.2 Å; H119, 2.2 Å; D145, 2.1 Å | none | Formally unassigned density (see **Fig. S5b**) | **Chain A:** S201 O–Fe 4.2 Å **Chain B:** S201 O–Fe 4.0 Å |

[a]Refinement statistics are presented in Table S1. [b]Bound at the positively charged pocket in the ST domain, very close to C408, invoking electrostatic steering. Close-ups of the Fe(II) coordination complexes are shown in Fig. S2. Hemifacial coordination of Fe(II) is by H56, H119, D145 in all cases. The number of modeled water molecules varies as indicated. If only one water is modeled, it occupies an axial coordination site hydrogen-bonded to H56.

**Table S3**. Comparison of EXAFS fits for plausible scatterer models of anaerobically prepared Fe-C408A CstB. The fit discussed in the main text (fit 10) is highlighted in **bold**.

| Fit | Scattering path | N[a] | R (Å) | $\sigma^2$ ($10^{-3}$ Å$^2$)[b] | $E_0$ (eV) | Red. $\chi^2$ | R-factor |
|---|---|---|---|---|---|---|---|
| 1 | Fe-(O/N)1 | 4 | 2.13 ± 0.01 | 4.2 ± 0.1 | 7123 ± 3 | 197 | 0.0889 |
| 2 | Fe-(O/N)1 | 5 | 2.13 ± 0.01 | 5.8 ± 0.1 | 7126 ± 3 | 194 | 0.0879 |
| 3 | Fe-(O/N)1 | 6 | 2.12 ± 0.02 | 7.3 ± 0.1 | 7121 ± 3 | 223 | 0.1010 |
| 4 | Fe-(O/N)1 | 1 | 2.00 ± 0.03 | 3.3 ± 3.0 | 7120 ± 2 | 107 | 0.0366 |
| | Fe-(O/N)2 | 3 | 2.13 ± 0.01 | 1.5 ± 0.1 | | | |
| 5 | Fe-(O/N)1 | 1 | 1.94 ± 0.03 | 2.0 ± 2.7 | 7117 ± 2 | 789 | 0.0269 |
| | Fe-(O/N)2 | 4 | 2.11 ± 0.01 | 2.4 ± 0.8 | | | |
| 6 | Fe-(O/N)1 | 1 | 1.90 ± 0.02 | 2.0 ± 2.3 | 7116 ± 2 | 87 | 0.0297 |
| | Fe-(O/N)2 | 5 | 2.10 ± 0.01 | 4.1 ± 0.7 | | | |
| 7 | Fe-(O/N)1 | 1 | 2.06 ± 0.03 | 2.4 ± 9.9 | 7122 ± 4 | 295 | 0.068 |
| | Fe-(O/N)2 | 3 | 2.17 ± 0.05 | 1.5 ± 6.8 | | | |
| | Fe-S1 | 1 | 3.27 ± 0.07 | 5.0 ± 6.8 | | | |
| 8 | Fe-(O/N)1 | 2 | 2.10 ± 0.04 | 3.4 ± 8.0 | 7122 ± 4 | 261 | 0.0610 |
| | Fe-(O/N)2 | 2 | 2.18 ± 0.09 | 1.5 ± 0.2 | | | |
| | Fe-S1 | 1 | 3.27 ± 0.07 | 5.5 ± 7.1 | | | |
| 9 | Fe-(O/N)1 | 1 | 2.00 ± 0.04 | 2.4 ± 3.3 | 7120 ± 3 | 206 | 0.0480 |
| | Fe-(O/N)2 | 4 | 2.15 ± 0.05 | 1.5 ± 0.1 | | | |
| | Fe-S1 | 1 | 3.25 ± 0.02 | 4.1 ± 4.7 | | | |
| **10** | **Fe-(O/N)1** | **1** | **1.95 ± 0.02** | **3.1 ± 1.9** | **7118 ± 1** | **22** | **0.0052** |
| | **Fe-(O/N)2** | **4** | **2.12 ± 0.02** | **2.6 ± 0.5** | | | |
| | **Fe-C1** | **3** | **3.12 ± 0.05** | **5.4 ± 1.9** | | | |
| 11 | Fe-(O/N)1 | 1 | 1.91 ± 0.01 | 2.7 ± 1.5 | 7117 ± 1 | 26 | 0.0059 |
| | Fe-(O/N)2 | 5 | 2.11 ± 0.01 | 4.2 ± 0.4 | | | |
| | Fe-C1 | 3 | 3.08 ± 0.02 | 5.0 ± 1.9 | | | |
| 12 | Fe-(O/N)1 | 1 | 2.07 ± 0.27 | 6.3 ± 29.3 | 7122 ± 4 | 283 | 0.0340 |
| | Fe-(O/N)2 | 4 | 2.14 ± 0.06 | 3.2 ± 3.7 | | | |
| | Fe-S1 | 1 | 3.40 ± 0.19 | 10.0 ± 0.2 | | | |
| | Fe-C1 | 3 | 3.12 ± 0.05 | 2.3 ± 4.5 | | | |

[a]N signifies scattering path degeneracy. [b]$\sigma^2$ is the Debye-Waller factor.

**Table S4.** Summary of detected species in both anaerobic and aerobic persulfidation assays using C408A CstB.[a]

| Observed Mass (Da) | Mass difference from the C408A CstB parent mass | Possible adduct(s) | Expected mass of this adduct(s) (– xH) |
|---|---|---|---|
| 50360 | 0 | Intact C408A CstB | 0 |
| 50392 | 32 | $O_2$ or S | 31 |
| 50408 | 48 | $O_3$ | 47 |
| 50422 | 62 | 2S or 4O | 63 |
| **50438** | **78** | **$SO_3$** | **79** |
| 50458 | 98 | 3S | 95 |
| 50507 | 147 | $SSSO_3$ | 143 |
| 50538 | 178 | 1 HPE-IAM | 177 |
| 50568 | 208 | 1S and 1 HPE-IAM | 209 |
| 50586 | 226 | 1 $O_3$ and 1 HPE-IAM | 224 |
| 50599 | 239 | 2S and 1 HPE-IAM | 241 |
| **50617** | **257** | **$SO_3$ and HPE-IAM** | **256** |
| **50647** | **287** | **$SSO_3$ and HPE-IAM** | **288** |
| 50714 | 354 | 2 HPE-IAM | 354 |
| 50746 | 386 | 1S and 2 HPE-IAM | 386 |
| 50778 | 418 | 2S and 2 HPE-IAM | 418 |

[a]Refers to ESI-MS spectra shown in Fig. 2d, *main text*. The entries that are in bold type are those adducts observed only in the presence of $O_2$.

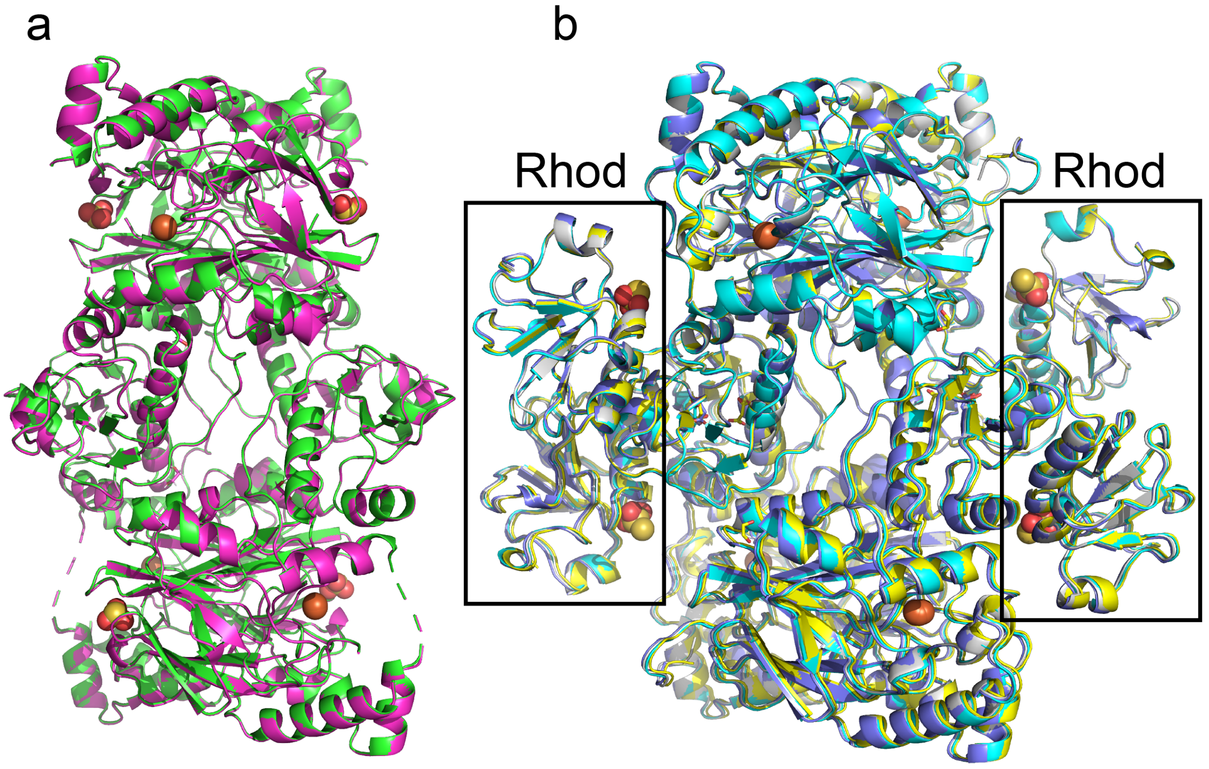


**Fig. S1.** Global structural alignmlovent of CstB variants. (**a**) Global structural alignment of two independent structures of wild-type CstB, in which the C201 harboring loop and the C-terminal Rhod domain are not visible. The structures in which C20 is persulfidated (PDB 28MT, *green*) and C20 is sulfonated (28MU, *magenta*) are shown. (**b**) Global structural alignment of wild-type CstB (28MV, *cyan*), C408S CstB (28MX, *yellow*), C201S/C408S CstB (28MY *gray*), and C201S CstB (28MW *purple*). See Table S1 for structure statistics for these models.

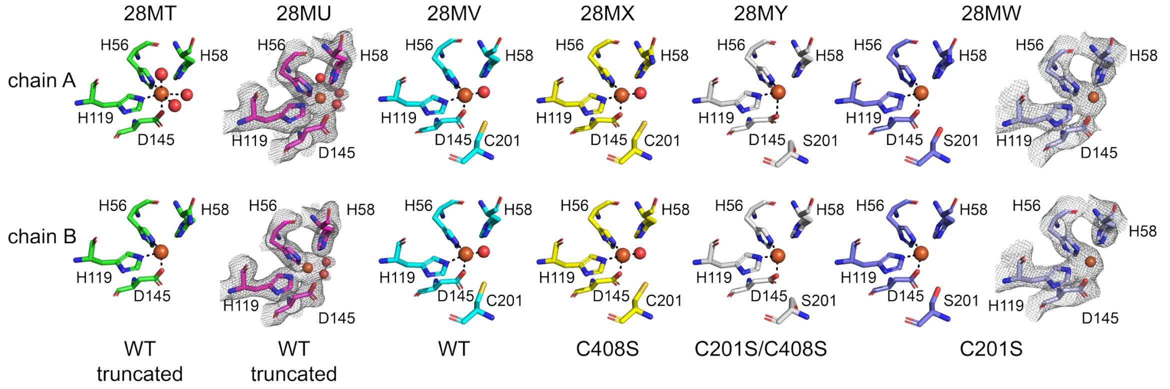


**Fig. S2.** Fe(II) active sites in each of the two chains for all six CstB structures reported here (see Table S1 for refinement statistics). Fe (shaded *rust*)-coordination bond distances and other information are provided in Table S2. Water molecules, *red* spheres. *Gray*, maps 2mFc - DFo contoured at σ=0.9 (28MU) and σ=0.9 (28MW).

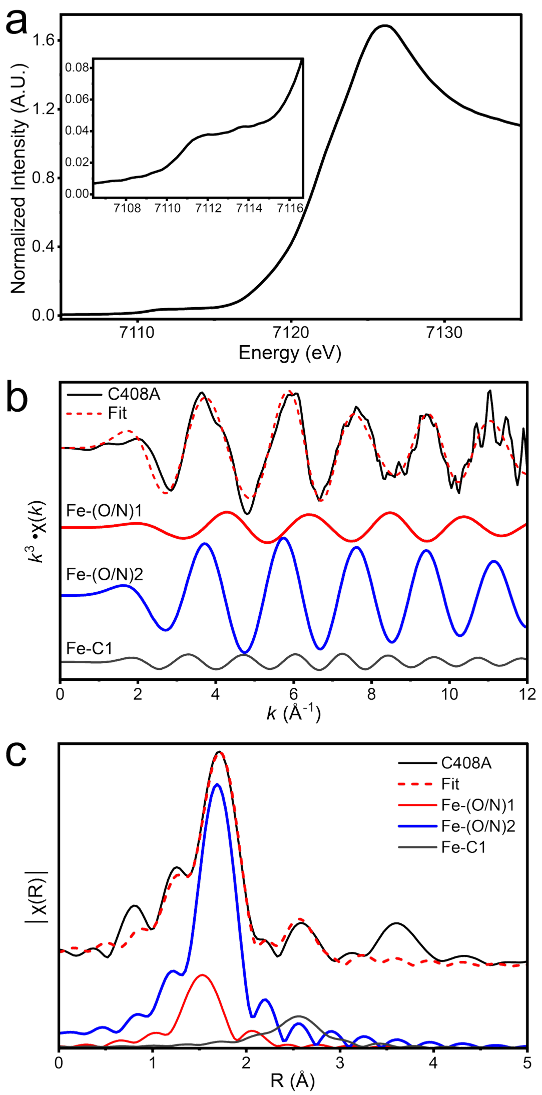


**Fig. S3. XAS data and fitting. (a)** Fe K-edge XAS spectra of C408A (*black*) showing normalized edge. Inset of the Fe K-edge XAS focuses on the pre-edge region of C408 from 7107 to 7117 eV. **(b)** EXAFS of C408 (*black*) in *k*-space with a fit corresponding to the best overall fit (dashed red line) and its individual components (colored lines) for C408A in Table S3. Spectra are $k^3$-weighted. **(c)** EXAFS of C408 (*black*) in R-space with a fit corresponding to the best overall fit (dashed *red* line) and its individual components (colored lines) for C408A in Table S3. Spectra are $k^3$-weighted, and FTs for spectra were performed across a $k$-range of 2-13 Å$^{-1}$. No phase shift has been applied.

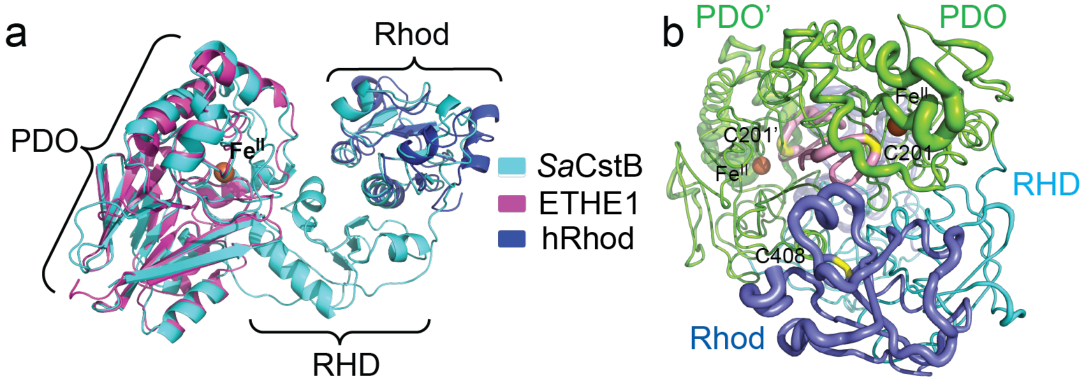


**Fig. S4. (a)** Global structural alignment of one subunit of the wild-type CstB (PDB ID 28MV, *cyan*), with human ETHE1 (PDB ID 4CHL, *purple*), and a human rhodanese, TSTD1 (PDB ID 6BEV, *blue*)[2,3]. (b) Sausage-plot of the domain-shaded wild-type CstB dimer, with the thickness of the cartoon corresponding to Cα B-factor. Plots like these for other mutant CstBs are largely identical to wild-type and thus are not shown here. The PDO domain is shaded *green* (Fe, *rust*), the RHD in *cyan* and the Rhod domain in *blue*. The C201 (shaded *yellow*)-containing active site loop is shaded *pink*.

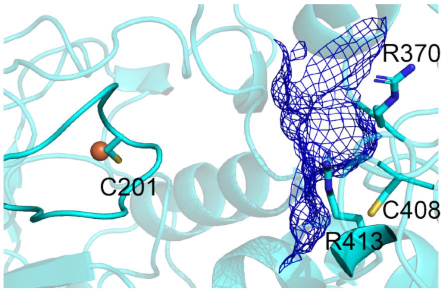


**Fig. S5.** Mesh representation of the surface of one subunit of the wild-type CstB reveals that R370 and R413 contribute to formation of a positively charged "wall" (*blue* mesh) that occludes the rhodanese active site represented by C201.

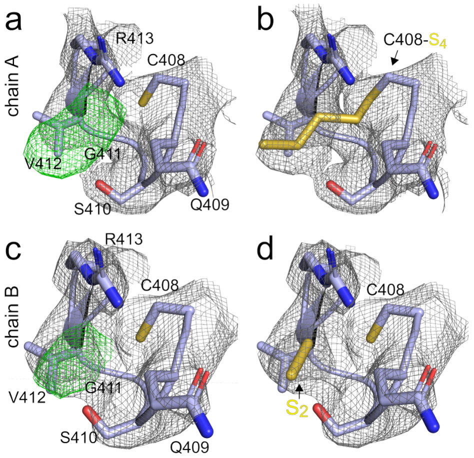


**Fig. S6.** Fitting of unassigned density (*green*) in the sulfurtransferase loop region around C408 in C201S CstB (28MW). (**a**) Chain A showing unassigned electron density (*green*, maps mFc – Dfo, contoured at σ=3.0). (**b**) Chain A unassigned electron density fitted with a polysulfide chain covalently bonded to C408 Sγ. (**c**) Chain B showing unassigned electron density (*green*, maps mFc – Dfo, contoured at σ=3.0). (**d**) Chain B unassigned electron density fitted with a molecule of $S_2^{2-}$ not covalently bonded to C408 Sγ. *Gray*, maps 2mFc - DFo contoured at σ=1.0 in all panels. Fitting this electron density with polysulfur chains, particularly in chain A is justified on the basis of the MS/MS data obtained for the wild-type enzyme (see Fig. S9b). The electron density in the deposited structure (28MW) is left unmodeled.

```
                                  130       140       150       160       170       180
S. aureus Newman       PESISFLLTDLGGGSSVPMGLFSGDFIFVGDIGRPDLLEKSVQIKGSTEISAKQMYESVQ
S. pasteuri            PESISFLLTDEGAGAQVPMGLFSGDFIFVGDIGRPDLLEKAVKVEGSSEIGAKQMFKSIK
S. epidermidis         PESISFLLTDEGAGAQVPMGLFSGDFIFVGDIGRPDLLEKAVKVEGSSEIGAKQMFKSIE
S. hominis             PESISFLLTDEGAGAQVPMGLFSGDFIFVGDIGRPDLLEKAVKVEGSSEIGAKQMFKSIE
S. equorum             PESISFLLTDEGGGSQVPMGLFSGDFIFVGDIGRPDLLEKAVQMEGSTEVSAEQMYHSIE
                       ********** *.*:.***********************:*:::**:*:.*:**: *::

                                  190       200       210       220       230       240
S. aureus Newman       NIKNLPDYVQIWPGHGAGSPCGKALGAIPISTIGYEKINNWAFNEIDETKFIESLTSNQP
S. pasteuri            SIKDLPDYIQIWPGHGAGSPCGKSLGAIPTSTLGYEKQTNWAFFENNEATFIDKLISDQP
S. epidermidis         SIKDLPNYIQIWPGHGAGSPCGKSLGAIPTSTLGYEKQTNWAFSENNEATFIDKLISDQP
S. hominis             SIKDLPDYIQIWPGHGAGSPCGKSLGAIPTSTLGYEKQTNWAFSENNEATFIDKLISDQP
S. equorum             TVKELSDSIQIWPGHGAGSPCGKSLGAVPMSTLGYEKMNNWAFNETNKVTFIETLTSNQP
                       .:*:*.: :**************:***:* **:**** .**** * ::..**:.* *:**
```

**Fig. S7.** The residues (highlighted in yellow) involved in van der Waals interaction to stabilize the loop are conserved in CstB proteins from other *Staphylococcus* strains.

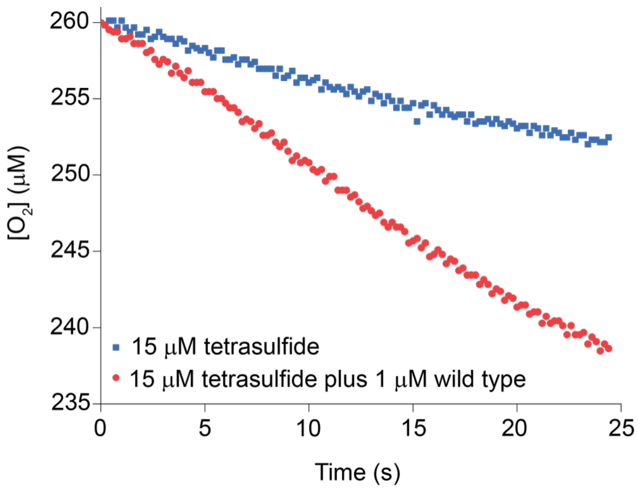


**Fig. S8.** Kinetic assay of the activity of wild-type *CstB* (1 μM) vs. sodium tetrasulfide (15 μM) as the persulfidation substrate.

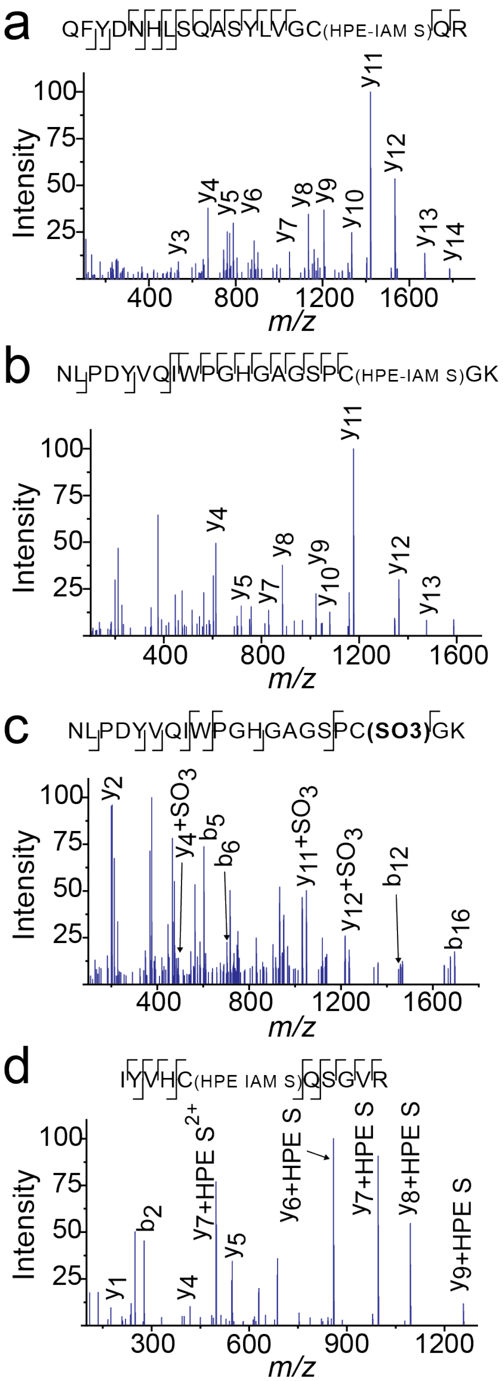


**Fig. S9.** MS-MS (MS2) analysis of the samples shown in Fig. 2d, main text. **(a)** Positive-ion mode MS2 spectrum of the peptide bearing a C201 that is persulfidated and capped with HPE-IAM. **(b)** Positive-ion mode MS2 spectrum of the tryptic peptide bearing a persulfidated and capped C408. **(c)** Positive-ion mode MS2 spectrum of the peptide bearing an *S*-sulfonated C201. **(d)** Positive-ion mode MS2 spectrum of the peptide bearing a persulfidated and capped C20.

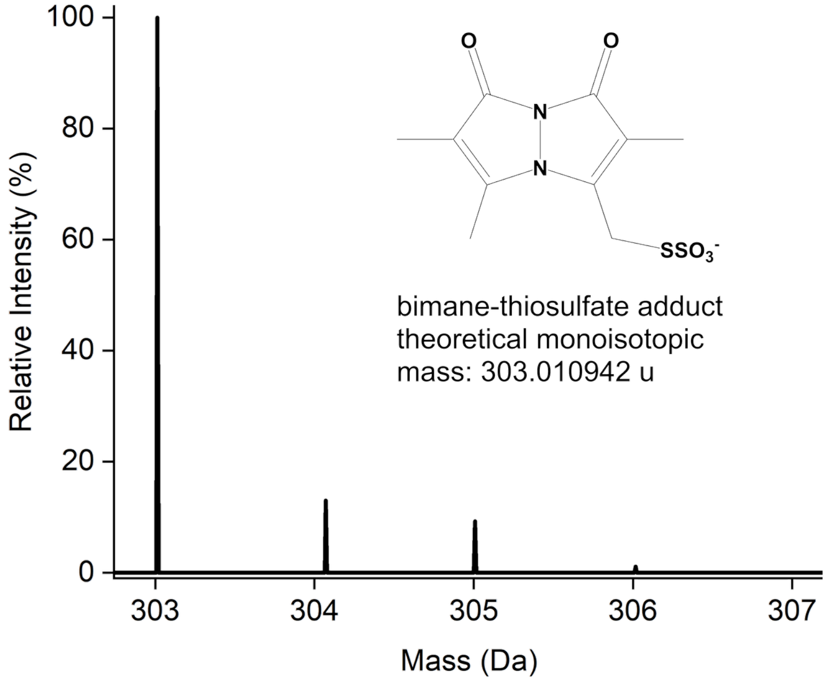


**Fig. S10.** Calculated normal isotope distribution for the isotopically "light" mBBr-TS adduct.

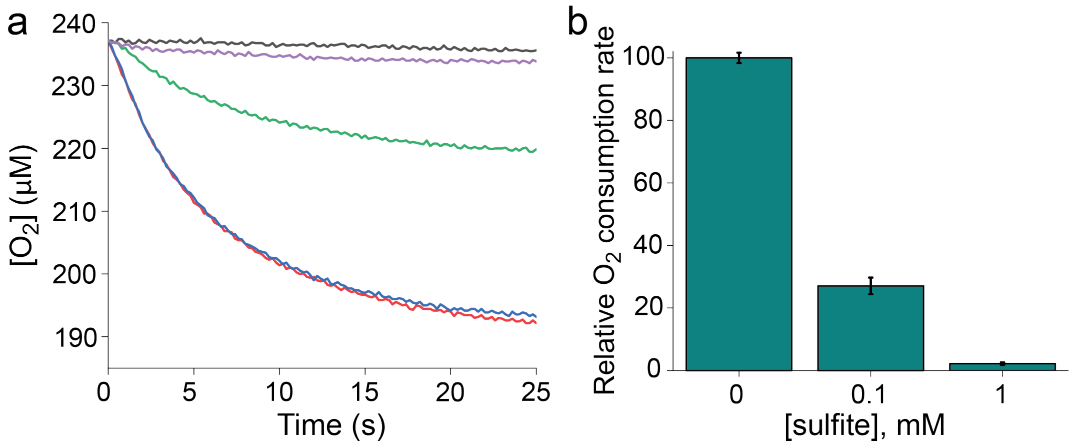


**Fig. S11.** Sulfite is an inhibitor of CstB oxygen consumption. **(a)** Representative $O_2$ consumption traces obtained for the 100 µM of CSSH only (*black* trace), WT CstB plus 100 µM of CSSH (*red*), WT CstB + 100 µM of CSSH + 1 mM EDTA (*blue*), WT CstB + 100 µM of CSSH + 1 mM EDTA + 0.1 mM sodium sulfite (*green*), and WT CstB + 100 µM of CSSH + 1 mM EDTA + 1.0 mM sodium sulfite (*violet*). **(b)** Summary of triplicate experiments like those shown in panel (a).

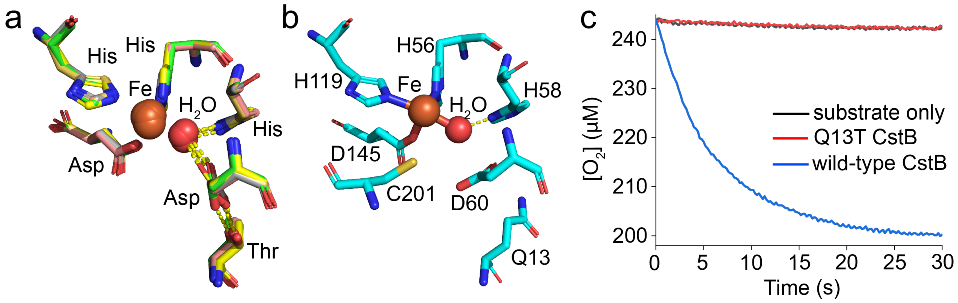


**Fig. S12. (a)** Fe active site crystal structure alignment of canonical PDOs, revealing the presence of a conserved Thr---Asp---$H_2O$ hydrogen-bonding network. **(b)** This H-bonding network is absent in the CstB Fe(II) active site, with Q13 replacing the Thr, giving rise to distinct orientation of H-bond donor D60. (c) $O_2$ consumption activity of Q13T CstB vs. the WT enzyme reveals that Q13T variant is inactive.

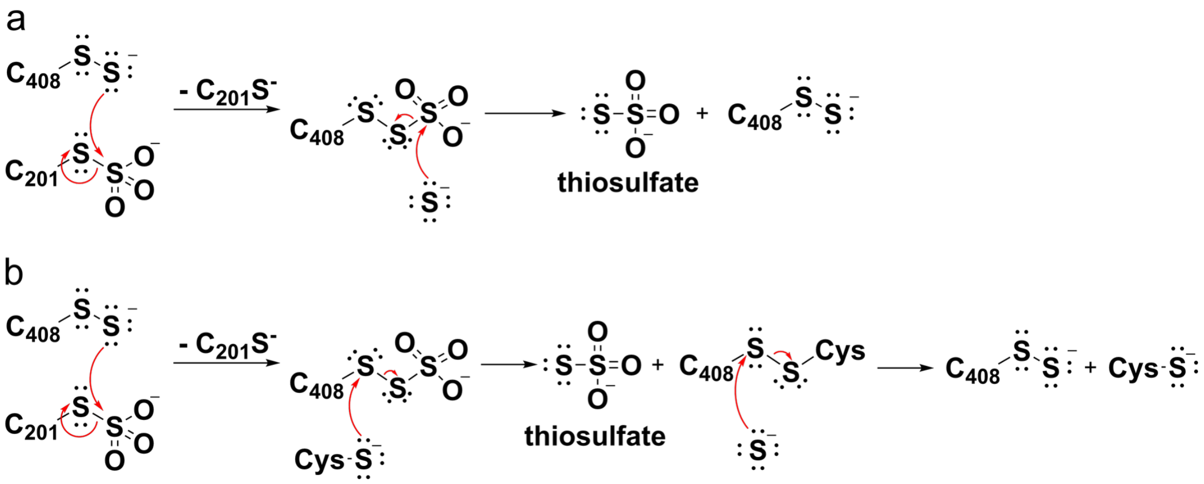


**Fig. S13.** Alternative routes to thiosulfate formation via canonical C201 *S*-sulfonate intermediate. (**a**) Contaminating sulfide from the substrate cocktail attacks the C408 *S*-sulfonate formed via *S*-sulfonate exchange with C201 *S*-sulfonate, to release thiosulfate, leading to preservation of the C408 persulfide. (**b**) The thiol of free cysteine product attacks to release thiosulfate and form a mixed disulfide, which contaminating sulfide then attacks C408 *S*-sulfonate to regenerate the C408 persulfide. Both mechanisms are inconsistent with the 2:1 RSSH:TS stoichiometry reported earlier[4], as the C408 cysteine persulfide persists following the initial turnover, thus consuming only one mol equivalent of RSSH after the first turnover.  Neither mechanism can be rigorously ruled on the basis of the work presented here.
